# Amazing structural diversity of giant virus-like particles in forest soil

**DOI:** 10.1101/2023.06.30.546935

**Authors:** Matthias G. Fischer, Ulrike Mersdorf, Jeffrey L. Blanchard

## Abstract

Large DNA viruses of the phylum *Nucleocytoviricota* infect diverse eukaryotic hosts from protists to humans, with profound consequences for aquatic and terrestrial ecosystems. While nucleocytoviruses are known to be highly diverse in metagenomes, knowledge of their capsid structures is restricted to a few characterized representatives. Here, we visualize giant virus-like particles (VLPs, diameter >0.2 µm) directly from the environment using transmission electron microscopy. We found that Harvard Forest soils contain a higher diversity of giant VLP morphotypes than all hitherto isolated giant viruses combined. These included VLPs with icosahedral capsid symmetry, ovoid shapes similar to pandoraviruses, and bacilliform shapes that may represent novel viruses. We discovered giant icosahedral capsids with structural modifications that had not been described before including tubular appendages, modified vertices, tails, and capsids consisting of multiple layers or internal channels. Many giant VLPs were covered with fibers of varying lengths, thicknesses, densities, and terminal structures. These findings imply that giant viruses employ a much wider array of capsid structures and mechanisms to interact with their host cells than is currently known. We also found diverse tailed bacteriophages and filamentous VLPs, as well as ultra-small cells. Our study offers a first glimpse of the vast diversity of unexplored viral structures in soil and reinforces the potential of transmission electron microscopy for fundamental discoveries in environmental microbiology.

## Introduction

Electron microscopy has been crucial for advancing the field of virology since Ruska and Knoll constructed the first “Übermikroskop” in the 1930s^1–4^. It allowed researchers to study the corpuscular nature of the previously “ultravisible” viruses and develop morphology-based virus classification systems that were popular until the 1970s^5, 6^. Twenty years later, electron microscopy was used to demonstrate that viruses were abundant and active in various aquatic environments^7, 8^, which helped us to recognize microbe-infecting viruses as key players in ecology and evolution^9–13^. Electron microscopy remains the method of choice for characterizing the particle structures and intracellular infection cycles of viruses, even though some extraordinarily large viruses can be seen with a light microscope.

These so-called “giant viruses” that infect mainly single-celled eukaryotes have particles in the size range of roughly 0.2 µm to 1.5 µm and carry DNA genomes that are up 2.5 million base pairs long^14–16^. Although large DNA viruses of unicellular algae had been studied for decades^17^, it was the discovery of Acanthamoeba polyphaga mimivirus that caused a conceptual breakthrough^18–20^. Mimivirus particles are covered with a dense layer of fibers that stained gram-positive and led to its misidentification as a coccoid bacterium, until electron microscopy revealed an icosahedral capsid shape typical of viruses^21^. With a capsid diameter of 0.5 µm and a genome of more than a megabase, mimivirus and many other giant viruses that were isolated in the following years blurred the earlier boundary between viruses and cells. As a practical consequence, the standard procedure of filtering water samples through 0.2 µm pore size filters to separate viruses from cellular microbes was thenceforth known to exclude an entire phylum of viruses. But how diverse are giant viruses in the environment? Culture-independent methods such as metagenomics have uncovered thousands of new giant virus genomes, proving that giant viruses are present in various aquatic and terrestrial ecosystems^22–24^. Analyses of metagenome-assembled genomes (MAGs) also facilitated an improved classification of giant viruses into several orders and families within the phylum *Nucleocytoviricota* (previously Nucleocytoplasmic Large DNA Viruses, NCLDVs)^25–28^, which includes mimiviruses (order *Imitervirales*), pandoraviruses and chloroviruses (order *Algavirales*), but also the poxviruses (order *Chitovirales*).

However, viral MAGs are often incomplete and contain little or no information about important biological traits such as host range, mode of infection, or structure and composition of the virus particle. These properties are best studied in virus-host systems that can be grown under controlled laboratory conditions. However, isolates of giant viruses are limited to a few different model systems, with a large bias towards amoebae-infecting viruses due to the successful use of *Acanthamoeba* spp. as a bait organism^29^. The fast pace of metagenomic sequencing combined with the slow progress in giant virus cultivation led to a growing discrepancy between a wealth of poorly understood viral MAGs and a shortage of well-characterized virus-host systems.

As a first attempt to alleviate this imbalance, we sought to complement metagenomic approaches with imaging methods and thus to investigate the morphological diversity of giant viruses directly from environmental samples. We chose a forest soil ecosystem for this study, as soils are known to harbor complex microbial communities, including diverse viral assemblages^30–32^. An estimated 97% of the ≈5E+31 virus particles on earth are found in sediments and soils^33^, but only some groups of soil-dwelling bacteriophages have been studied in greater detail^34^. One of the few studies that have focused on giant viruses in soil used fluorescence-activated cell sorting to reduce the complexity of cells and viruses in samples from an experimental soil warming experiment at Harvard Forest. This mini-metagenomics approach revealed 16 new MAGs of NCLDVs, including a member of the *Mimiviridae* with a 2.4 megabase pair genome^23^. The discovery of these giant virus genomes in Harvard Forest soil inspired us to study the morphological diversity of virus particles from the same location.

In this work, we show that natural soil virus assemblages can be characterized by high-quality negative staining transmission electron microscopy (TEM) to reveal ultrastructural details of uncultured viruses. We report a high diversity of giant virus-like particles in the 0.22 µm to 1.2 µm size fraction and describe previously unknown viral morphotypes. Of particular interest are icosahedral particles with unique structural modifications such as double capsids, channels, portals, tails, fibers, and other types of appendages. We also describe a variety of bacteriophage morphologies and soil nanoparticles that resembled pandoraviruses, filamentous viruses and ultra-small prokaryotes. Our study provides a visual counterpart to the metagenomic diversity of soil viruses, which is known to be higher than in most aquatic environments^32^.

## Materials and Methods

### Origin and processing of soil samples

Soil cores were collected on Aug 28, 2019 from a site adjacent to the Barre Woods (coordinates 42.48, −72.18) long-term warming experiment where a previous metagenomics study on giant viruses was conducted^23^ and at a site adjacent to the Prospect Hill (coordinates 42.54, −72.18) long-term warming experiment at the Harvard Forest in Petersham, MA, USA. The soil and site characteristics have been previously detailed^35, 36^. In brief, the soils are coarse-loamy inceptisols in mixed hardwood forest stands with the dominant tree species being paper and black birch (*Betula papyrifera* and *B. lenta*), red maple (*Acer rubrum*), black and red oak (*Quercus velutina* and *rubra*), and American beech (*Fagus grandifolia*). A clear demarcation of the organic (approximately top 5 cm of soil core) and mineral (below 5 cm of soil core) layers allowed for manual separation. The soil samples were subsequently stored at 4°C at the University of Massachusetts Amherst. After obtaining approval on Sep 2, 2019 from the Regierungspräsidium Karlsruhe (Plant Protection Service) under community directive 2008/61/EG, a subset of the samples was shipped to the Max Planck Institute for Medical Research in Heidelberg, Germany.

Five hundred milliliters of autoclaved and 0.22 µm filtered mineral water (Volvic, Danone Germany GmbH) were added to 100 g of soil and mixed by shaking and stirring for 15 min. The suspension was passed through a colander to remove roots and stones and then centrifuged for 5 min at 500x g, 20°C. The supernatant was filtered sequentially through 47 mm diameter filters (Millipore, Germany) with 20 µm (NY-20), 5 µm (TMTP) and 1.2 µm (RTTP) pore sizes. Material retained on these filters was discarded and the 1.2 µm filtrate was further passed through a 0.22 µm (GTTP) filter. The 0.22 µm filter was transferred to a sterile 50 ml plastic tube so that the retentate side was facing inwards. Particles were recovered from the filter by adding 5 ml of sterile mineral water and rocking the tube for 2 h, followed by storage at 4°C. Particles in the resulting suspension were concentrated by centrifugation at 100,000x g for 30 min, 18°C using an SW40 rotor in an Optima XE-90 ultracentrifuge (Beckman, Germany). The supernatant was discarded and the pellet was resuspended in approximately 50 µl of remaining supernatant.

### Electron microscopy

The resuspended pellet was mixed with 0.5% paraformaldehyde and incubated at 20-25°C for 30 min. Ten microliters of the fixed virus sample were pipetted onto a formvar/carbon-coated and glow-discharged copper EM grid and incubated for 10 min. Supernatant from the grid was removed by blotting with filter paper and the grid was washed and blotted three times with double-distilled water. A drop of 0.5% uranyl acetate was added to the grid and incubated for 30 sec, then the grid was blotted and dried. Samples were examined using an FEI Tecnai G2 T20 TWIN transmission electron microscope (FEI, Eindhoven, The Netherlands) operating at 200 kV accelerating voltage. Electron micrographs were recorded with an FEI Eagle 4k HS, 200 kV CCD camera. For best possible resolution and contrast, we performed an optimized basic alignment and a daily routine alignment of the electron microscope. Focus settings and astigmatism corrections were optimized for each image.

### Image processing

TEM images were opened in Adobe Photoshop 24.2.1 and scaled to the desired dimensions (Image size menu) at their native resolution of 96 dpi using the “Preserve details 2.0, 0% noise reduction” resampling option. The resized image was then copied into the composite image file (at 300 dpi) and levels were adjusted individually for each image of the figure. Scale bars were retraced and formatted manually, and the automatically generated scale bars were cropped from the final images. The tailed VLP in Fig. 3j is a composite of two individual images for higher resolution. Particle measurements were made in ImageJ V1.53t.

### Criteria for classifying virus-like particles (VLPs)

In this study, we investigated the diversity of VLPs in forest soil with a focus on VLPs that share similar features with giant viruses of protists. We emphasize that it is not possible to prove the viral nature of any particles analyzed only by negative staining TEM, regardless of how similar to published virus structures they may look. However, we can assign different levels of confidence to VLPs from environmental samples, based on previously described specimens of viruses, microorganisms and other nanoparticles (Suppl. Fig. 1). Among the most recognizable virus structures are head-and-tail bacteriophages and accordingly, we are confident that VLPs with isometric or prolate heads and thin, tubular tails probably correspond to viruses of the class *Caudoviricetes*. Gene transfer agents (viriforms) have appearances similar to siphoviruses, but their head diameter is typically below 40 nm and thus smaller than most tailed bacteriophages^37–39^.

Many virus particles have icosahedral base symmetry, which results in planar projections of regular hexagons or penta-/decagons. Such virus particles often display 5- or 6-fold symmetry, depending on the type of virus, its orientation on the EM grid, and the effects of staining and dehydration during sample preparation. We are therefore quite confident that particles with pentagonal or hexagonal cross sections represent viral capsids, especially when regular surface patterns indicate the presence of capsomers. However, we acknowledge that capsid-like structures such as encapsulins or carboxysomes can be released from cells and may be mistaken for VLPs in electron micrographs^40–42^. Furthermore, we cannot exclude that some very large capsids with icosahedral symmetry may deviate from the canonical capsomer-based architecture and could lack repeating surface structures. Larger VLPs may also be more prone to damage and deformation during sample preparation than smaller VLPs, which can impede their classification.

We were generally more cautious when interpreting the nature of filamentous, ovoid and bacilliform particles, even when they are composed of repeating subunits. While many of them are likely to represent virions, it is difficult to distinguish between a filamentous virus and detached cellular structures or broken bacteriophage tails. Similarly, pandoraviruses, pithoviruses, and other eukaryotic viruses with large non-isometric particles are probably found in these soil samples, but to categorize all of them as viruses would include false positives, such as small prokaryotic cells with regular surface structures. Pandora- and pithoviruses have been mistaken for cellular microorganisms in the past^43–45^, and the paucity of isolates for these virus groups makes it difficult to delineate clear morphological identification criteria.

At the bottom of our confidence scale are particles in the 50-2000 nm size range that cannot be attributed to known viral or cellular morphologies. Some of these may represent new viral morphotypes of damaged viral capsids, but many are probably non-viral entities such as small cells, vesicles, organic scales, cell fragments, and inorganic particles. Furthermore, we do not make claims about specific hosts for any VLPs, although we assume that VLPs with caudoviricete-like morphologies infect bacteria or archaea, and that large (>200 nm) VLPs with icosahedral or ovoid morphology are most likely to infect eukaryotic hosts.

### Reference samples

Negative staining TEM is a simple yet effective method for studying ultrastructural features such as the shape, surface and size of virus particles. However, particles can shrink as a result of dehydration during sample preparation. To better compare environmental VLPs with cultured viruses, and because the sample preparation and image quality of published electron micrographs of giant viruses vary considerably, we analyzed representatives of pithoviruses, molliviruses, pandoraviruses, and marseilleviruses (all courtesy of L. Bertaux and C. Abergel, Univ. Aix-Marseille), as well as Cafeteria roenbergensis virus (CroV) and its virophage mavirus using the same negative staining TEM protocol that we applied to the soil samples.

## Results

### Particle structures of reference protist viruses

To best compare soil VLPs with previously described giant virus structures, we recorded reference images for six different giant viruses and one virophage (Fig. 1). Our selection of viral isolates includes some of the largest and smallest known protist viruses (compare Fig. 1a and g) and thus serves as a good starting point for exploring the morphological diversity of giant viruses in environmental samples. *Pithovirus sibericum* particles are ovoid particles with a reported length of up to 2.5 µm^46^. We measured lengths between 900 nm and 1550 nm, and widths between 550 nm and 720 nm. The apical pore with the striated cork structure was visible in several particles (Fig. 1a, top). Some particles were covered with a 20 nm thick fiber layer. Overall, these measurements are within the range of previous reports, although they are slightly smaller than pithovirus particles analyzed by Cryo-EM^47^. *Pandoravirus neocaledonia*^48^ particles were ovoid with a length of 925-1135 nm and a width of 660-785 nm. Compared to the dimensions reported for other pandoraviruses (0.8-1.2 µm x 0.5 µm)^14^, *P. neocaledonia* virions appeared to be slightly wider. The unique apical pore, also called ostiole, was clearly visible in most particles (Fig. 1b) and serves as a distinguishing feature for this group of viruses. The roughly spherical virions of *Mollivirus sibericum* are 500-600 nm in diameter and covered with a hairy tegument^49^. The particles we analyzed were slightly larger with an average diameter of 681±28 nm and an external fiber layer that was 60-85 nm thick (Fig. 1c). Depending on the orientation of the virions, some particles were slightly ovoid with a length of 720-750 nm and a width of 560-620 nm.

**Figure 1.**
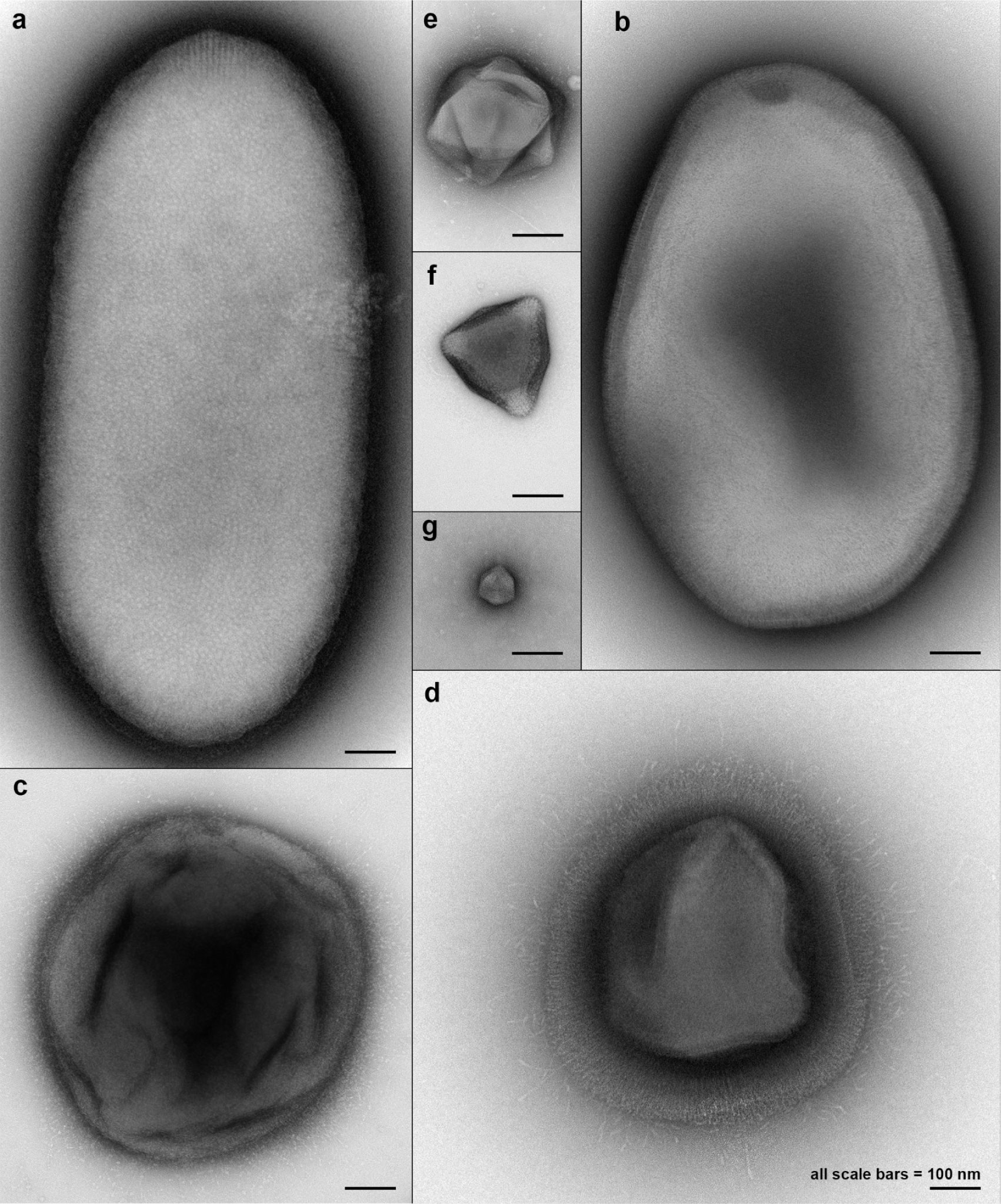
Reference negative staining electron micrographs of cultured protist viruses. **a)** *Pithovirus sibericum*. **b)** *Pandoravirus neocaledonia*. **c)** *Mollivirus sibericum*. **d)** Acanthamoeba polyphaga mimivirus. **e)** Cafeteria roenbergensis virus. **f)** Melbournevirus. **g)** The virophage mavirus. All scale bars, 100 nm.

Acanthamoeba polyphaga mimivirus has capsids with icosahedral base symmetry that were reported to be 400 nm (TEM^21^) to 500 nm (cryo-EM^50^) in diameter. One vertex is a modified genome exit portal named the stargate^51^, which is covered by a protein structure called the starfish complex^52^. With our negative staining protocol, we measured the average vertex-to-vertex diameter of mimivirus capsids to be 459±33 nm. The starfish structure was visible in many, but not all particles (Fig. 1d, top). On the exterior, mimivirus capsids are covered with a dense layer of fibrils that have been reported to be up to 125 nm long^50, 53^. Here, we noticed that most virions featured two distinct fiber layers, an inner, denser one with an average fiber length of 84±18 nm, and an outer, less dense layer with single fibers protruding for an additional distance of 100±21 nm (Figs. 1d, 4a). The first publication of CroV mentioned hexagonal particles that were 230-300 nm in diameter^54^, and later cryo-EM studies reported 300 nm-wide particles^55^. For this study, we measured an average capsid diameter of 288±12 nm (Fig. 1e). Melbournevirus was isolated from a freshwater pond in Melbourne, Australia and had 200 nm-wide icosahedral particles in thin-section TEM^56^ and ≈230 nm-wide capsids in cryo-EM^57^. We measured an average capsid diameter of 220±11 nm (Fig. 1f). Capsids of the virophage mavirus were regular icosahedrons with an average vertex-to-vertex diameter of 76±4 nm (Fig. 1g), which matches previous reports^58^.

### The Harvard Forest micro-gallery

We then analyzed VLPs in the 0.22-1.2 µm size fraction of organic and mineral layer soil from two locations at Harvard Forest in Petersham, MA, USA, near long-term experimental warming plots. We collected 684 TEM images with a preference for giant VLPs, including 426 images from Prospect Hill and 258 images from Barre Woods. Of these images, 565 originated from the organic soil horizon, and 119 from the mineral zone (Suppl. Fig. 2a). Overall, the diversity of VLPs and cell-like particles was much higher in the organic horizon than in the mineral layer, which explains the larger dataset for the former. In addition, 12 images were taken of a pooled concentrate from the size fraction smaller than 0.22 µm. We classified ≈350 particles as VLPs, including ≈300 isometric, ≈30 ovoid and ≈20 filamentous VLPs. About 110 images with cells or cell-like particles were collected and the remaining ≈220 images showed particles that we could not confidently categorize (Suppl. Fig. 2b).

### Virus-like particles with icosahedral symmetry

We found icosahedral particles in a wide range of sizes (Fig. 2). The majority of VLPs in this category appeared to be plain capsids that lacked identifiable surface modifications. We note, however, that many particles were surrounded by fibers of varying nature that could not be clearly identified as capsid-attached structures. In addition, fragile capsid appendages may have become detached during sample handling.

**Figure 2.**
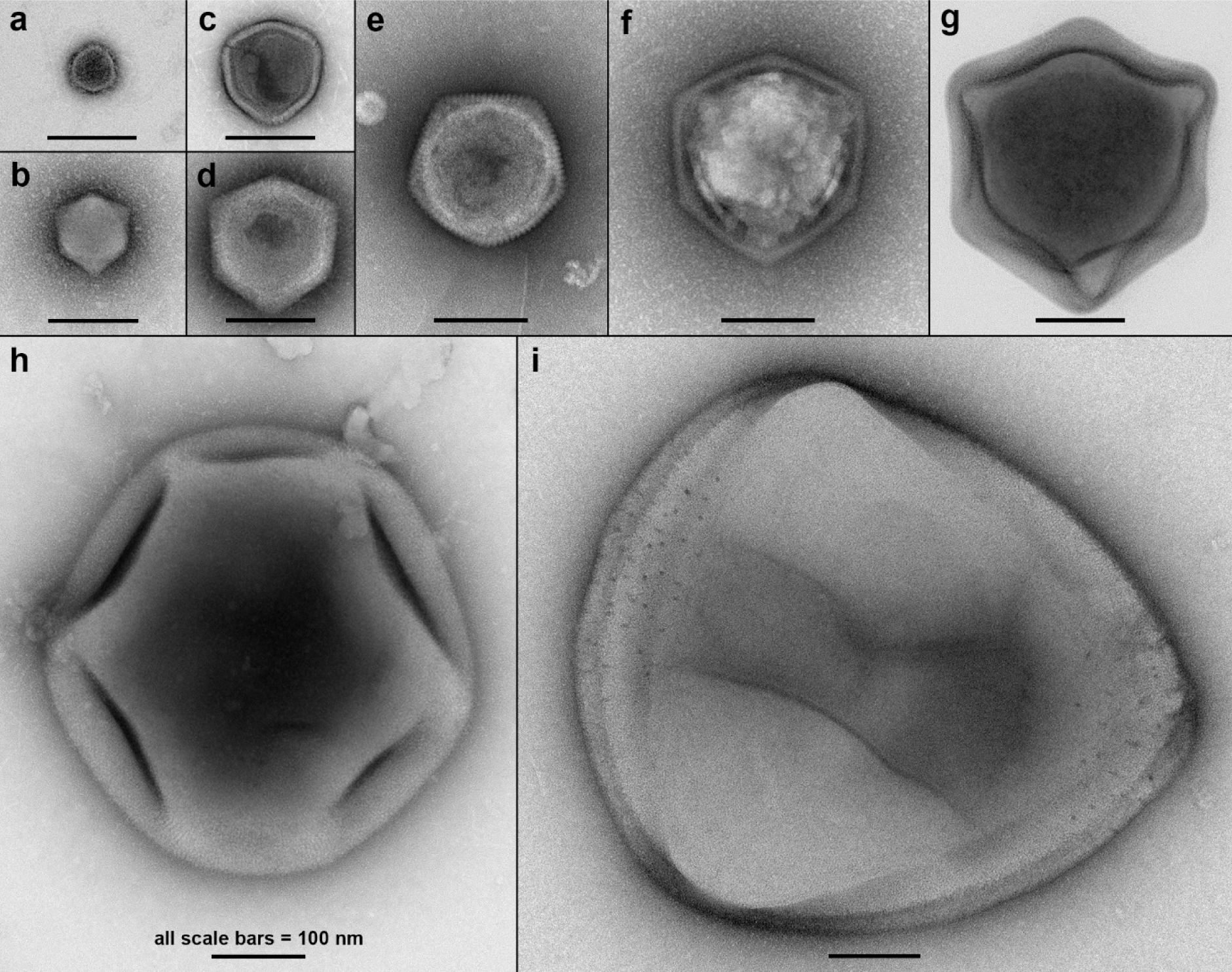
Virus-like particles with plain icosahedral capsids. Shown is a selection of VLPs with capsid diameters ranging from 50 nm to 635 nm, which corresponds roughly to a 2000-fold difference in volume. All scale bars, 100 nm.

The average vertex-to-vertex diameter of plain icosahedral capsids ranged from 50 nm to 635 nm. The smallest of these were in the size range of virophage capsids (50-90 nm, Fig. 2a,b), while others resembled marseilleviruses or CroV (Fig. 2f,g). We observed an even distribution in the 50-350 nm size range, whereas plain icosahedral capsids larger than 350 nm were relatively rare (Suppl. Fig. 2c). The actual size distribution may be even more skewed towards smaller particles, because many VLPs smaller than 200 nm may have passed through the 0.22 µm filters from which we recovered the VLPs in this study.

Curiously, we found that several giant VLPs with isometric or round shapes had distinct clusters of more darkly stained spots with pore-like appearance (Suppl. Fig. 3). These pore-like structures were on average 5.6±0.7 nm in diameter and were sometimes arranged in groups of 3 to 7 spots in one particular area of the capsid (Suppl. Fig. 3a-n), whereas other VLPs had more widely distributed spot patterns (Fig. 2i, Suppl. Fig. 3o+p). We did not observe VLPs that were smaller than 200 nm in diameter and had such spots. These features may not necessarily correspond to pores or holes in the capsid layer, caused for instance by missing capsomers or depressions, but may simply reflect a greater binding affinity for uranyl acetate of the proteins located there, which leads to positive staining. The proteins or capsomers in these locations could potentially serve as anchor points for internal virion structures such as scaffolding proteins or lipid membranes.

### Giant VLPs with previously unseen capsid modifications

To our surprise, we found that many capsids displayed structural modifications that had not been described before. Isometric VLPs with a diameter of >200 nm often featured tails, modified vertices, double capsid layers, and tubular appendages or internal structures (Fig. 3, Suppl. Figs. 4,5). Of these new morphotypes, one resembled the general appearance of mimivirus, with a ≈430 nm wide capsid that was covered in a dense layer of 100-150 nm fibers (Fig. 3a). In contrast to mimivirus, however, no stargate portal was visible and some giant VLPs seemed to consist of two nested capsid structures, the inner one with a diameter of ≈350 nm (Fig. 3a). Other particles that lacked the additional outer capsid layer had a diameter of 315-370 nm with an 80-100 nm thick layer of fibers (Suppl. Fig. 4a+b).

**Figure 3.**
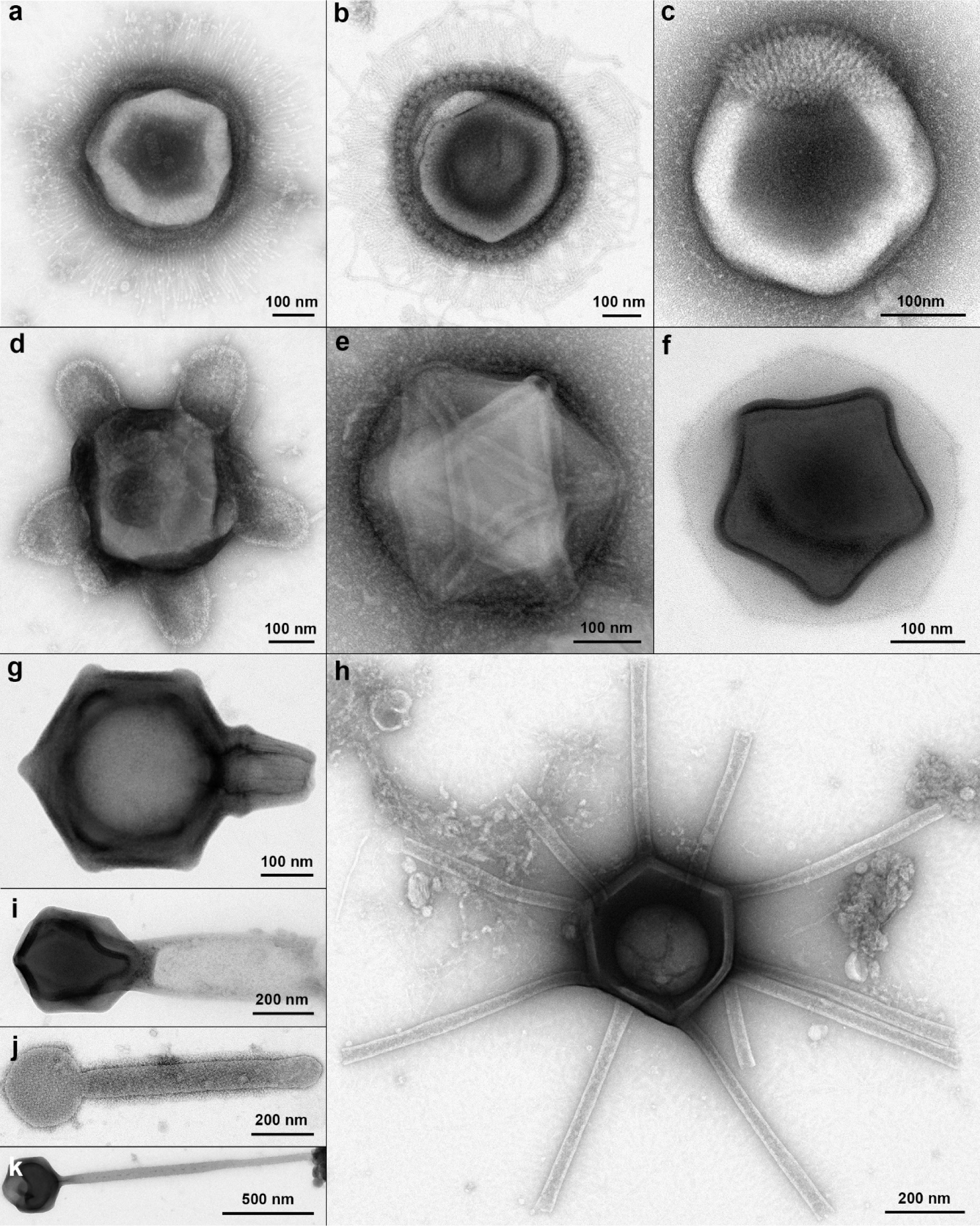
Morphotypes of giant virus-like particles with unique structural features from Harvard Forest soil. **a)** “Mimi-like” morphotype. **b)** “Supernova” morphotype. **c)** “Haircut” morphotype. **d)** “Turtle” morphotype. **e)** “Plumber” morphotype. **f)** “Christmas star” morphotype. **g)** “Flacon” morphotype. **h)** “Gorgon” morphotype. **i)**-**k)** Large VLPs with tail structures.

The related yet distinct “supernova” morphotype had a spherical outer capsid layer with a diameter of 490 nm, and an isometric-looking inner capsid with a diameter of 380 nm (Fig. 3b). The outer capsid layer was ≈35 nm thick and consisted of regularly arranged multimeric capsomers that were ≈23 nm wide. Capsomers of the inner capsid were smaller and more tightly arranged. External ≈135 nm long fibers were associated with the outer capsomers. We recorded similar VLPs of lower quality that had outer capsid diameters of 400 nm and 485 nm (Suppl. Figs. 4c+d). Inside the cores of some particles, we observed one or two coiled cylindrical structures, which were 15-25 nm wide, up to 80 nm long, and appeared to be helical with a periodicity of 8-11 nm (Fig. 3a, Suppl. Figs. 3k, 4c). Although these structures are much smaller than the helical genomic fibers of mimivirus^59^, they could also contain viral nucleic acid.

The “haircut” category comprised several VLPs with 280-325 nm large capsids and asymmetrically distributed outer fibers of varying lengths, thicknesses and densities (Fig. 3c, Suppl. Fig. 4g-i). The “turtle” morphotype was characterized by lobed appendages that protruded from the ≈380 nm wide capsids (Fig. 3d, Suppl. Fig. 4e). Each lobe was ≈150 nm long and ≈140 nm wide and consisted of regularly arranged subunits. The attachment points and number of lobes per particle are not clear, but we hypothesize that each lobe extends from a capsid vertex. Modified vertices were also found in the “plumber” morphotype, but these structures penetrated into the interior of the 340-400 nm wide capsids (Fig. 3e, Suppl. Fig. 4f).

Each vertex contained a pore structure with an inner diameter of 15-20 nm, creating a channel through the capsid layer. In one instance, the vertices were connected with each other by a network of intra-capsid channels (Fig. 3e).

One giant VLP morphotype that we found repeatedly in Harvard Forest soil was the “Christmas star”, which consisted of a double-layered shell where an electron-dense inner capsid with a diameter of 285 nm was nested within a 385 nm wide and less electron-dense outer capsid (Fig. 3f, Suppl. Fig. 4j-l). The vertices of the inner capsid always aligned with the faces of the outer capsid, and the inner capsid had a 10-13 nm thick wall structure. However, the thickest capsid walls reported to date for an icosahedral VLP were found in the “flacon” morphotype. Its 420-470 nm wide capsids were fortified with a 35-70 nm thick outer wall, and all except one of the vertices were capped with a rounded and slightly elevated structure (Fig. 3g, Suppl. Fig. 5g+h). The remaining vertex was transformed into an ≈150 nm long conical nozzle that was ≈160 nm wide at its base and ended into an opening with a 25-50 nm inner diameter (Suppl. Fig. 5i). One “flacon” VLP had a more cylindrically shaped vertex extension, with a thin thread of material attached to it (Suppl. Fig. 5h+j). This structure may represent the genome exit portal of the virion.

Among the most unusual giant VLPs was the “Gorgon” morphotype with its long tubular appendages (Fig. 3h, Suppl. Fig. 5a-f). The capsids had a diameter of 410±20 nm with a ≈20 nm thick wall. Each particle had 8-11 tubular extensions, which were 500-650 nm long and 30-65 nm wide. The appendages were straight, presumably hollow with an opening at the distal end, and composed of regularly arranged subunits with a periodicity of ≈4.5 nm (Suppl. Fig. 5e). At the proximal end, they were attached to a capsid vertex (Suppl. Fig. 5f). We assume that each vertex had one appendage, with the possible exception of a unique vertex for genome release.

Lastly, we found several 260-450 nm wide VLPs with diverse tail structures. Some tails were long and thick (Fig. 3i: 560×210 nm; Suppl. Fig. 5k: 660 x 160 nm), some were covered with fibers (Fig. 3j: 750×90 nm tail, 25 nm long fibers), and another one was 1.4 µm long and 30-50 nm wide (Fig. 3k).

### Giant VLPs are covered with diverse fiber structures

We then analyzed the diversity of fibrous structures on giant VLPs in more detail. Capsid-associated fibers among NCLDVs can be short and sporadic such as in PBCV-1^60^, or dense as on the 260 nm-wide capsid of Medusavirus, an *Acanthamoeba*-infecting giant virus, which is covered with 14 nm long fibers that have a spherical end structure^61^. The long and dense fiber layer of mimivirus consists of glycoslyated proteins and is thought to mimic bacterial peptidoglycan and also to increase the virion diameter in order to trick their amoebal hosts into phagocytosing them. Mimivirus fibers are 120-140 nm long, 1.4 nm wide, and terminate in globular structures at the distal end^62–64^. In our negative staining TEM analysis, mimivirus fibers presented as two distinct layers, a denser one close to the capsid wall with a thickness of 60-110 nm, and a thinner outer one with single fibers extending ≈120 nm above the first layer (Figs. 1d, 4a). Notably, fibers in both layers terminated in small globular heads.

**Figure 4.**
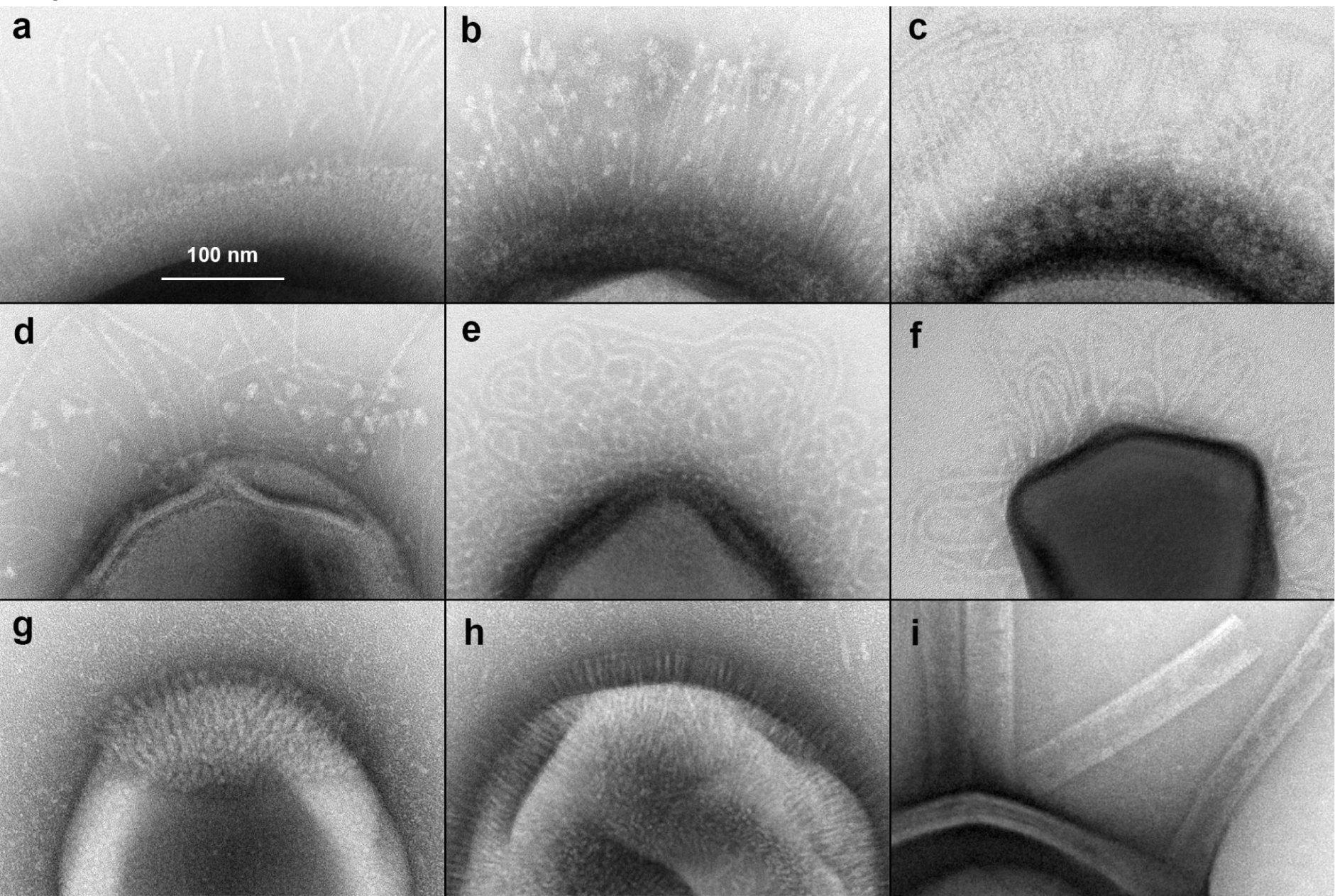
Diversity of capsid fibers among large virus-like particles with icosahedral symmetry. **a)** Mimivirus reference. **b)** “Mimi-like” morphotype. **c)** “Supernova” morphotype. **d)** Long thin fibers with triangular end structures. **e)** Long thin curly fibers with globular end structures. **f)** Long thin fibers without apparent terminal structures. **g)** “Haircut” morphotype with short, thin fibers partially covering the capsid. **h)** VLP with short, thin fibers covering the entire capsid. **i)** “Gorgon” morphotype with long and thick appendages. The scale bar in a) applies to all images.

In the Harvard Forest samples, we observed giant VLPs with diverse types of fibers attached to the capsid surfaces (Fig. 4). In the “mimi-like” morphotype, the external fibers were not as dense as in mimivirus, ≈120 nm long and ≈4 nm wide and capped with ≈7 nm wide globular structures (Fig. 4b). The fibers surrounding capsids of the “supernova” morphotype were ≈115 nm long and 6 nm wide and also had globular tips with a diameter of 7-8 nm (Fig. 4c). We found one VLP that was covered with a sparse layer of disordered, 100-200 nm long and 3 nm thin fibers with ≈12 nm wide triangular heads (Fig. 4d). Another VLP had a dense, 100-200 nm thick layer of curly, ≈4 nm wide fibers whose individual lengths were difficult to measure but which ended in small globular tips (Fig. 4e). Several VLPs featured a few dozen ≈4 nm wide and ≈700 nm long fibers without any apparent end structures (Fig. 4f).

The “haircut” category comprises different morphotypes with asymmetrically attached fibers, such as the ≈4 nm wide and ≈18 nm long fibers that covered ≈15% of the VLP in Figs. 3c+4g, or the ≈120 nm long bundle of ≈3 nm wide fibers shown in Suppl. Fig. 4h. Other particles had peritrichous fibers that were ≈26 nm long and ≈2.5 nm wide (Fig. 4h). In contrast to the above examples, the “Gorgon” appendages are tubular structures that extend from the capsid vertices and are likely to serve a different function (Figs. 3h, 4i, Suppl. Fig. 5a-f).

### Giant VLPs with ovoid morphology

Although many viruses of the phylum *Nucleocytoviricota* have capsids with icosahedral symmetry, ovoid morphotypes are common among poxviruses, ascoviruses, pandoraviruses, and pithoviruses. We found a variety of ovoid nanoparticles in Harvard Forest soil (Fig. 5), and although we considered only those with regular surface patterns as giant VLPs, we cannot exclude that some of them may be of cellular nature. Pandora- and pitho-/cedratviruses possess unique features that can help to identify them as VLPs, namely an apical pore and one or two apical cork structures, respectively (Fig. 1a+b).

**Figure 5.**
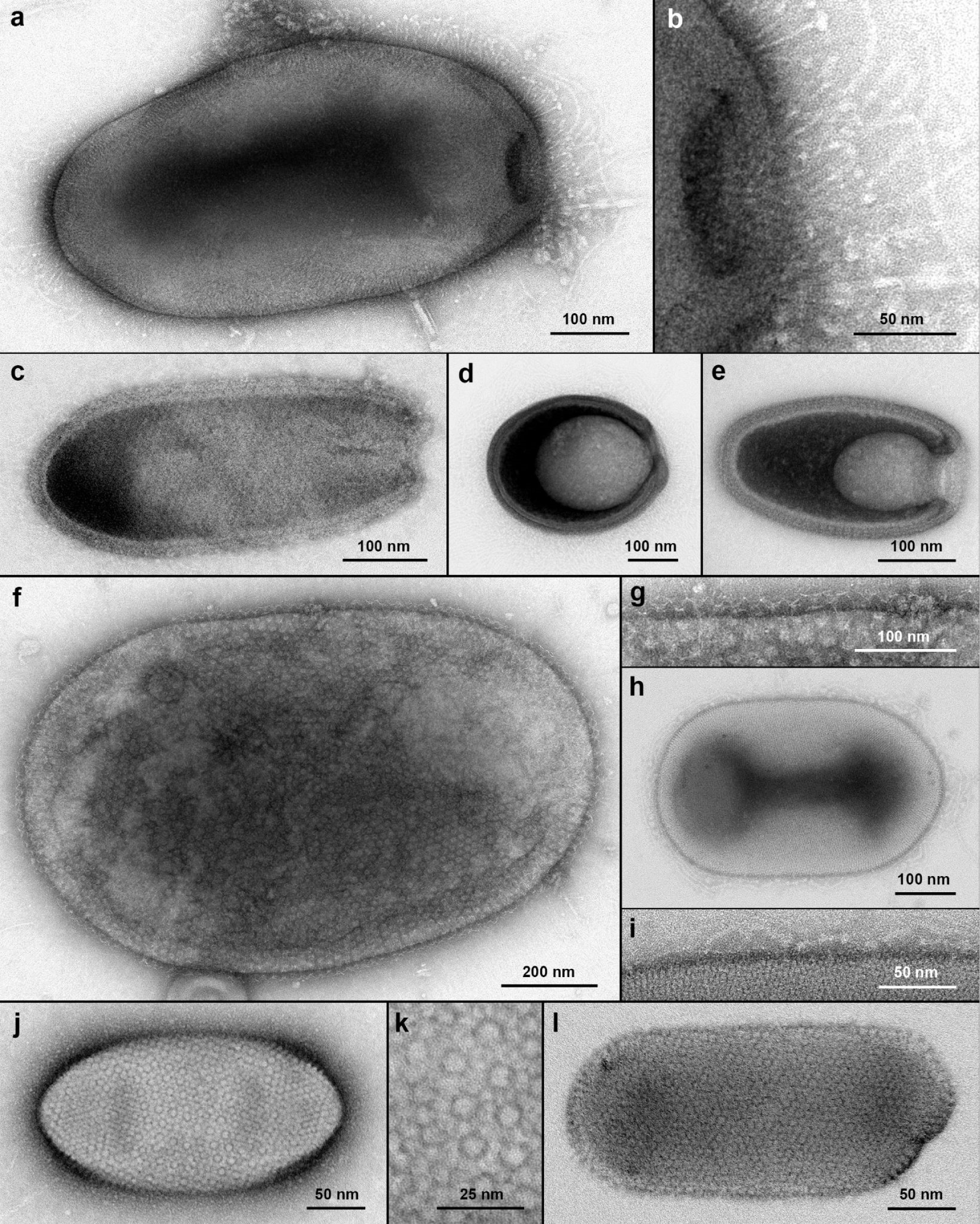
Ovoid particles from Harvard Forest soil. **a)** 630×330 nm particle with a ≈80 nm wide pore, partially covered in ≈40 nm long fibers with globular tips. A few longer fibers protrude from the pore apex. **b)** High magnification view of the VLP in a). **c)** 460×200 nm particle with regular surface pattern and a ≈50 nm wide pore. **d)** 355×275 nm particle with a ≈60 nm wide pore, covered in a dense layer of 10-15 nm long fibers and a less dense layer of 50-200 nm long fibers. **e)** 310×180 nm particle with a 75 nm wide pore connected to an internal cavity that penetrates ≈135 nm into the particle. **f)** 1170×730 nm particle with ≈11 nm large surface subunits, covered with ≈15 nm long Y-shaped antennae. **g)** High magnification view of the particle in f). **h)** 470×290 nm particle with regularly arranged, ≈4 nm wide subunits. **i)** High magnification view of the particle in h). Single ≈4 nm wide fibers are visible on the surface. **j)** 265×140 nm particle with regular surface pattern. **k)** High magnification view of the VLP in j). Capsomer-like structures have a diameter of ≈6 nm and an average center-to-center distance of 13 nm. **l)** 280×125 nm particle with ≈5 nm spaced capsomer-like subunits.

In our dataset, several VLPs displayed apical pores (Fig. 5a-e), although none of them were as big as pandoravirus isolates. The largest ones were 530-630 nm long and ≈330 nm wide, with a pore diameter of 60-80 nm (Fig. 5a). By comparison, the *P. neocaledonia* pore structures measured 60-75 nm in diameter. Unlike pandoraviruses, these soil VLPs were unevenly covered with ≈2 nm thin and ≈40 nm long fibers that had 4-6 nm wide globular heads (Fig. 5b). Other VLPs in this category were 310-460 nm long and 180-275 nm wide with pore diameters of 50-75 nm (Fig. 5c-e). We did not find any particles with cork structures, but some VLPs in the 1 µm range had regular surface patterns that were reminiscent of pithoviruses (Fig. 5f+g).

### Filamentous particles

Rod-shaped and filamentous virus forms are associated with hosts across all domains of life. For instance, plant viruses of the family *Alphaflexiviridae* have flexible filamentous virions that are 400-800 nm long and 10-15 nm in diameter. Bacterial viruses of the family *Inoviridae* have flexible particles that are 6-10 nm wide and 600-2500 nm long. A variety of different filamentous virions can be found among archaeal viruses, such as the 600-900 nm long and 23 nm wide stiff rods of rudiviruses, the 410-2200 nm long and 24-38 nm wide flexible lipothrixvirus filaments, the enveloped 400×32 nm rods of tristromaviruses, or the rigid 143×16 nm rods of clavaviruses^65^. Many filamentous viruses of prokaryotes have unique structures at their particle ends, such as caps, fibers, spikes, or claws that provide additional identification criteria.

We found several filamentous particles in Harvard Forest soils that probably depict virions (Suppl. Fig. 6), including stiff rods with lengths of 380-600 nm and diameters of 20-50 nm (Suppl. Fig. 6a-e) or long flexible VLPs (Suppl. Fig. 6f). Many of them were composed of regularly spaced subunits suggestive of a helical architecture. Other forms of previously undescribed filamentous particles with head structures were also present (Suppl. Fig. 6g).

### Tailed bacteriophages

Bacteria constitute the most abundant soil microorganisms; accordingly, tailed bacteriophages were the most prominent group of VLPs we encountered in the 0.22-1.2 µm soil fractions. However, we recorded only a subset of these as we were mainly interested in giant VLPs. In total, we imaged 65 putative myoviruses with long and contractile tails (Suppl. Fig. 7), 27 siphoviruses with long and non-contractile tails (Suppl. Fig. 8), and 22 podoviruses with short and non-contractile tails (Suppl. Fig. 9). The largest of these may represent jumbo bacteriophages, which are defined by genome lengths exceeding 200 kbp and particles with head diameters larger than about 100 nm and tail lengths of approximately 100-500 nm^66^. Heads of myo-like VLPs were mostly isometric and had diameters of 55-150 nm (99±25 nm, Suppl. Fig. 2d)), those of sipho-like VLPs were either isometric (n=15) with diameters of 51-143 nm (79±27 nm) or prolate (n=12) with lengths of 77-392 nm and widths of 30-70 nm, and heads of podo-like VLPs were isometric with diameters of 59-116 nm (73±15 nm).

In the literature^67^, myovirus tails are described as being 80-455 nm long and 16-20 nm thick; siphovirus tails as being 65-570 nm long and 7-10 nm thick, and podovirus tails as being about 20 nm long and 8 nm thick. In the Harvard Forest soil samples, we identified myo-like VLPs with tails that were 63-234 nm (126±36 nm) long (Suppl. Fig. 2d) and 14-27 nm (20±4 nm, excluding contracted tails) thick. Most tails of sipho-like VLPs were 67-551 nm (245±106 nm) long and 6-19 nm (11±3 nm) thick, with the notable exception of a single siphovirus that had a 1.05 µm long tail (Suppl. Fig. 8b). Tails of podo-like VLPs were 9-67 nm (21±13 nm) long and 5-31 nm (13±5 nm) thick.

### Potentially novel viral morphotypes

At the bottom of our VLP confidence scale (Suppl. Fig. 1) were soil nanoparticles for which we could find no similar examples in the literature, but which could nevertheless represent virions. However, these may also represent cellular microbes, cell fragments, or other non-viral entities. A selection of such structures is shown in Fig. 6, including spherical particles with nested layers (Fig. 6a) or external fibers (Fig. 6b), and particles consisting of two subunits that were connected by a zipper-like structure with 9 nm long teeth (Fig. 6c). We also found droplet- and ovoid-shaped particles that lacked apparent capsomer-like surface patterns and often showed differently stained features along the longitudinal axis (Fig. 6d-f), and dome-shaped particles in the 400-700 nm size range (Fig. 6g-i). Lastly, we noticed a couple of 200-650 nm wide particles that resembled equilateral triangles with rounded corners and symmetrically arranged internal features (Fig. 6j-l).

**Figure 6:**
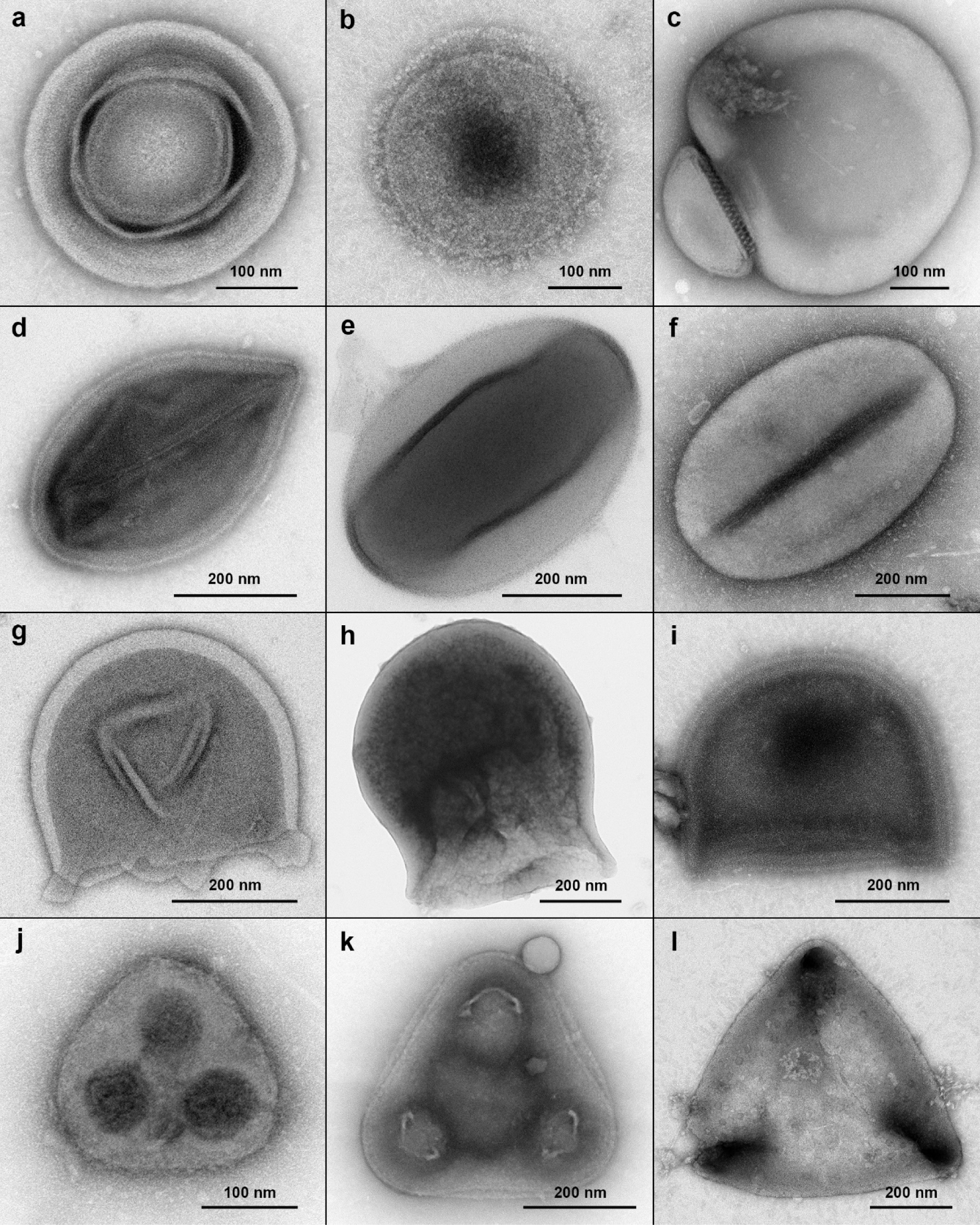
Unclassified soil nanoparticles. A selection of particles with round, ovoid, dome-shaped, and triangular morphologies that may represent new viral morphotypes, ultrasmall microorganisms, or subcellular structures. **a)** Multi-layered particle with a 330 nm outer diameter 330 nm and an inner sphere diameter of 180 nm. **b)** Vesicular particle with a diameter of 280 nm and 20-35 nm long external fibers with globular heads. **c)** Particle consisting of a larger 470×430 nm subunit and a smaller 240×100 nm subunit. **d)** Droplet-shaped particle, 500×310 nm. **e)** Ovoid particle, 565×380 nm. **f)** Ovoid particle, 635×415 nm. **g)** Dome-shaped particle with a 390 nm diameter. **h)** Dome-shaped particle, 690×580 nm. **i)** Dome-shaped particle, 480×390 nm. **j)** Triangular particle with a corner-to-corner distance of 200 nm. **k)** Triangular particle with a corner-to-corner distance of 425 nm. **l)** Triangular particle with a corner-to-corner distance of 650 nm.

### Cellular microorganisms

We imaged multiple particles with prokaryote-like appearances, although it was not possible to distinguish between bacteria and archaea in negative staining TEM (Suppl. Fig. 10). Some of these cells had VLPs attached to their surfaces (Suppl. Fig. 11). Due to our pre-filtration procedure, most cell-like particles were below 1 µm in at least one dimension. The smallest such objects measured 100×200 nm with a cytoplasmic volume of 0.001 µm^3^ and were surrounded by a ≈50 nm thick extracellular matrix (Suppl. Fig. 12). Their morphological features are in agreement with ultra-small bacteria of the Candidate Phyla Radiation (CPR), a diverse clade of mostly uncultured bacteria that have small streamlined genomes and presumably symbiotic lifestyles^68, 69^. Metagenomic signatures of CPR bacteria are found in Harvard Forest soil, including sequences matching the TM6 group and the phylum *Patescibacteria*^70^ (Blanchard, unpublished). We also cannot exclude that some of the tiny cells in our dataset may be DPANN archaea^71^. No obvious cases of protist cells were found and we assume that most soil protists were removed by the 1.2 µm pore-size filtration step.

## Discussion

Transmission electron microscopy of organic and mineral layer soil from Harvard Forest revealed an astounding diversity of virus-like particles. Of particular interest are the various appendages and other modifications of large icosahedral capsid structures: these include tubular protrusions, fibers of different lengths and thicknesses, internal channels, double capsids, unique vertex structures, and tails. Although electron microscopy alone is not sufficient to unambiguously establish the nature of an observed nanoparticle, we argue that large icosahedral VLPs with modified capsids can be called “virus particles” with even higher confidence than some of the most iconic virus morphotypes: the head-tail morphotypes typical of bacteriophages (Suppl. Fig. 1). This is because gene transfer agents may be visually mistaken for tailed bacteriophages, whereas we are not aware of any non-viral particles that would resemble large, appendage-bearing capsids with icosahedral-based symmetry.

Amazingly, we found that a few hundred grams of forest soil contained a greater diversity of capsid morphotypes than that of all hitherto isolated giant viruses combined. This observation is even more astounding when considering that we imaged only an infinitesimally small fraction of the viral diversity present in these soil samples. Viral abundances in forest soils have been reported to range from 10^8^ to more than 10^9^ VLPs per gram dry weight^32^. Most virus particles were probably removed by our filtration procedure, because virions are known to adhere to mineral and organic soil particles^72^. Viral recovery of at least certain groups of viruses can be improved by the use of specialized buffers, density gradient fractionation, or more rigorous mechanical treatment^73, 74^. Our goal was to obtain high-quality images of intact VLPs, rather than to maximize viral yields at the cost of introducing additional bias or damaging VLPs during the extraction procedure; hence, we opted for the simpler method of suspending the soil in sterile mineral water with minimal shaking. For this reason, and because we preferentially recorded giant VLPs while ignoring many tailed bacteriophages, we abstain from drawing quantitative conclusions about the viral morphotypes we found in Harvard Forest soil. However, given the enormous diversity of soil viruses^32^, the fact that we encountered some giant virus morphotypes multiple times suggests that they were abundant in these samples (e.g., “Christmas star” and “Gorgon” morphotypes, Suppl. Figs. 4+5).

Metagenome analysis of soil from the same location resulted in genome assemblies for 16 novel giant viruses^23^. These included relatives of pithoviruses, tupanviruses, and klosneuviruses, for which we found potentially matching VLP types: large ovoid, tailed icosahedral, and plain or fibered icosahedral particles, respectively. Although we currently cannot link metagenome-assembled genomes to any of the morphotypes described here, we show that the morphological diversity of giant VLPs clearly exceeds the metagenomic diversity of giant viruses in Harvard Forest soil.

With this visual display of viral diversity, we hope to inspire other researchers to explore different microcosms by electron microscopy, and to isolate more virus-host systems for detailed characterizations. New model systems are needed to elucidate the function and evolutionary origin of capsid appendages in giant viruses. Some structures may be analogous or homologous to cellular counterparts. For instance, the tubular appendages of the “Gorgon” morphotype resemble extracellular tubules of the hyperthermophilic archaeon *Pyrodictium abyssi*^75^.

Our study raises the question whether the extraordinary diversity of viral morphotypes is more typical of soil ecosystems, or whether appendage-bearing giant VLPs are also common in aquatic environments. Due to a lack of electron microscopy-based environmental surveys, especially for size fractions larger than 0.2 µm^30, 76^, this issue remains unresolved for now. For instance, a TEM study on marine water samples collected during the Tara Oceans Expedition reported that 50-90% of VLPs were tailless with an average diameter of 50 nm, whereas the remaining VLPs resembled tailed bacteriophages^77^. In contrast, large icosahedral capsids with tail structures were found in the Southern Ocean^78, 79^, the North Pacific Ocean^80^, and the North Sea^81^, suggesting that giant viruses with tails and other capsid modifications may be widespread in oceanic environments.

We discovered an unexpected diversity of soil VLPs in the 0.2 µm to 1.2 µm size fraction, which is typically excluded from virome studies. The cornucopia of viral morphotypes found in Harvard Forest alone questions our current understanding of the virosphere and its structural heterogeneity. This fascinating window into the complex world of soil viruses leaves little doubt that the high genetic diversity of giant viruses is matched by diverse and previously unimaginable particle structures, whose origins and functions remain to be studied.

## Acknowledgements

This work is dedicated to the memory of Hans-Wolfgang Ackermann (1936-2017), a pioneer in electron microscopy and taxonomy of bacteriophages. We think he would have enjoyed this display of viral diversity and hope that the image quality would have found his approval^82^. We are grateful to Lionel Bertaux and Chantal Abergel (Univ. Aix-Marseille) for providing pure samples of amoebal giant viruses. We thank Reinhard Rachel for valuable feedback and Rob Lavigne, Andrew Kropinski and Alexander Probst, and many others also, for discussions. Special thanks go to Karsten Richter (DKFZ Heidelberg) for sharing his expertise and passion for electron microscopy. The Max Planck Society enabled this study through funding and infrastructure support.

## Author contributions

**Matthias G. Fischer**: Conceptualization, Formal analysis, Investigation, Resources, Writing – Original draft, Writing – Reviewing and Editing, Visualization, Supervision. **Ulrike Mersdorf**: Methodology, Investigation, Formal analysis, Writing – Reviewing and Editing. **Jeffrey L. Blanchard**: Conceptualization, Formal analysis, Investigation, Writing – Reviewing and Editing.

## Supplementary Figures

**Supplementary Figure 1:**
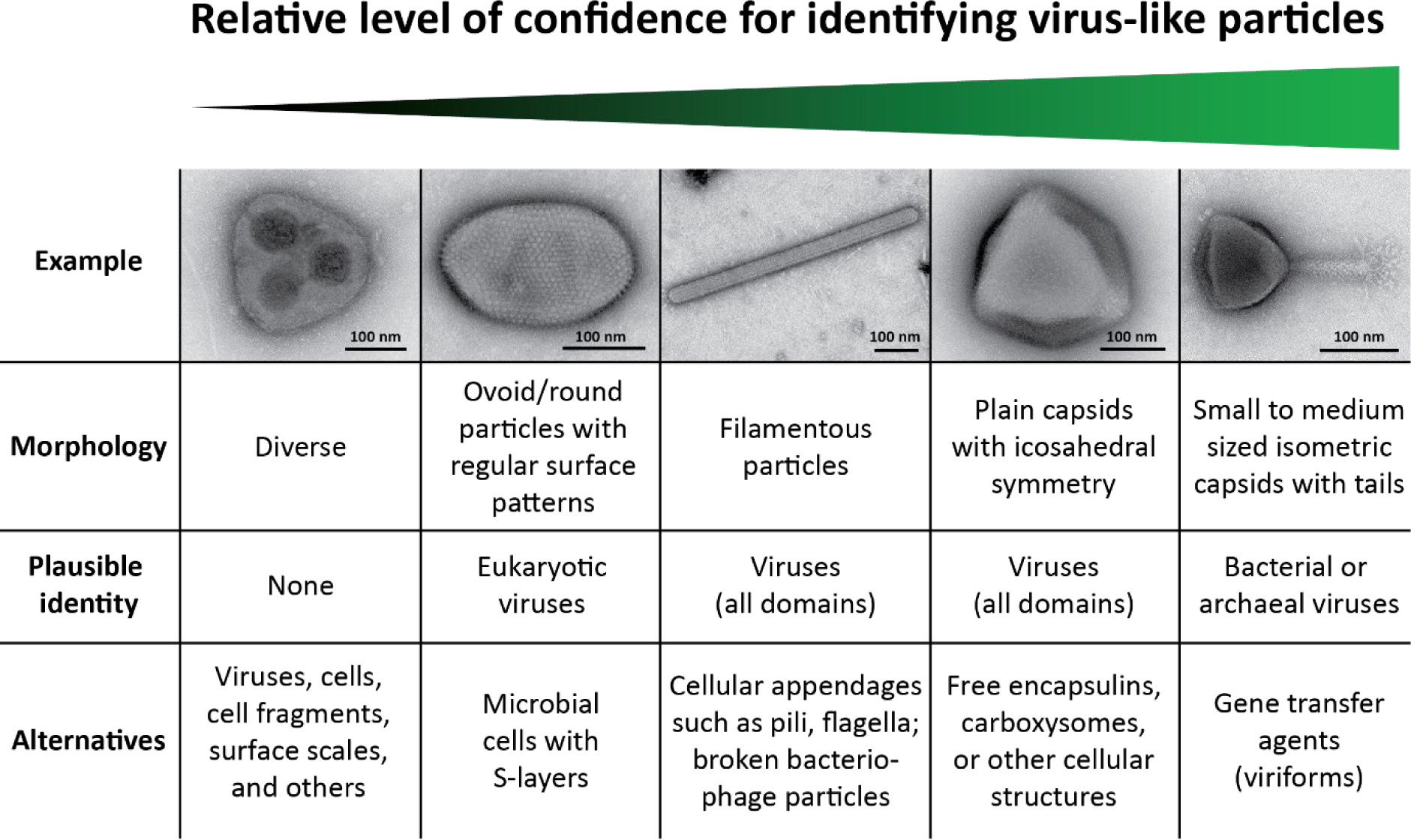
Considerations for classifying nanoparticles in the 50-2000 nm size range as VLPs. Identification of potential virus particles in TEM data is not trivial and depends on several factors including size, shape, and ultrastructure of the particle and its similarity to known structures, image quality, knowledge of technical limitations and potential artifacts, and experience and background of the researcher. The figure summarizes different categories of virus-related nanoparticles and their possible interpretations.

**Supplementary Figure 2:**
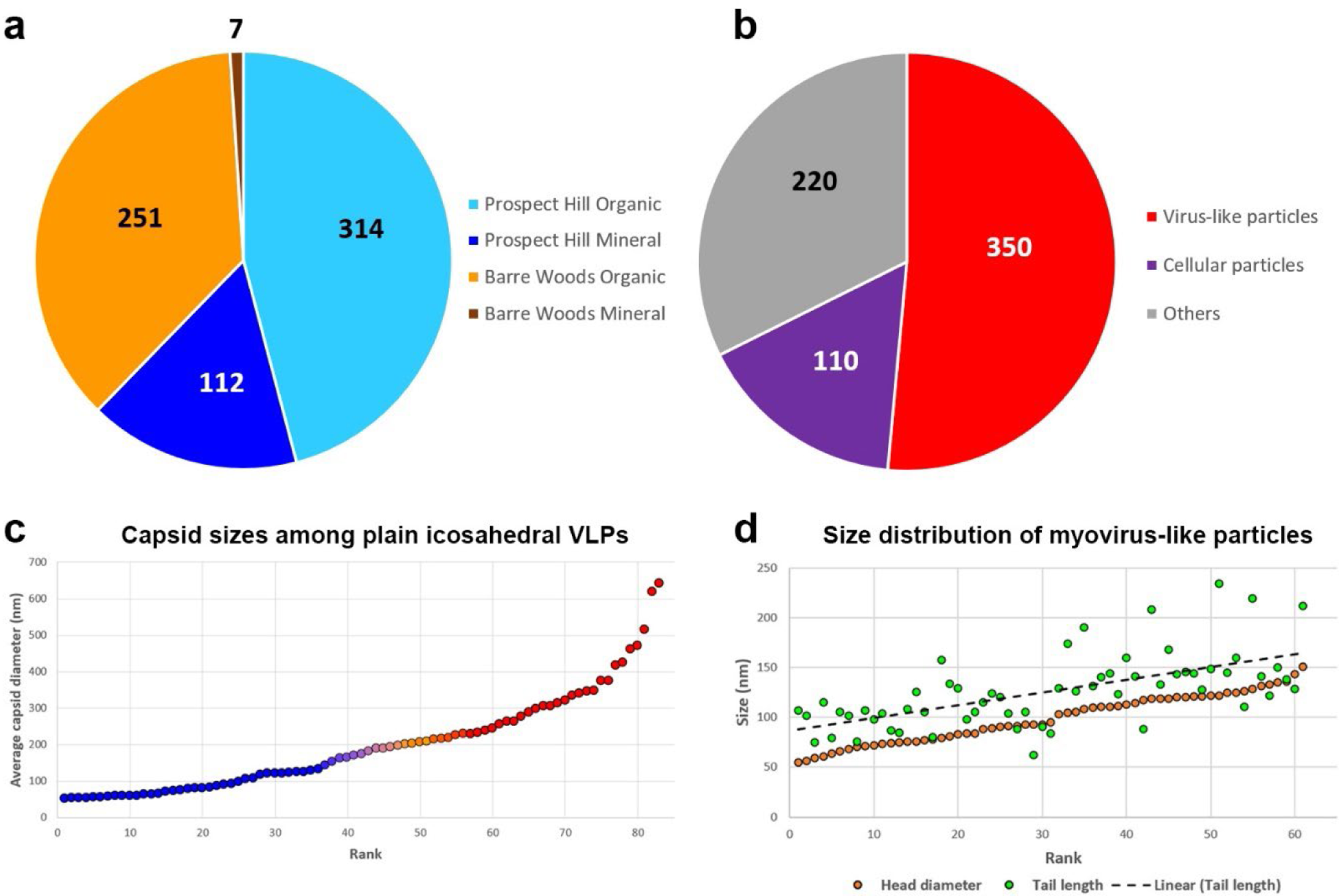
Electron microscopy data overview, capsid diameters, and myovirus head-tail measurements. **a)** Numbers of TEM images per sample. **b)** Numbers of TEM images by category. **c)** Distribution of capsid diameters among VLPs with plain icosahedral appearance. Capsid measurements smaller than 200 nm are shown in blue, those corresponding to giant VLPs are shown in red. As there is no clear size definition for giant viruses, the transition zone is colored as a gradient. **d)** Size distribution of tailed myovirus-like particles (A1 morphotype). Capsid diameters (orange) and tail lengths (green) are displayed separately. A linear regression of tail lengths shows that myovirus tails are on average 20-40 nm longer than their capsids are wide.

**Supplementary Figure 3:**
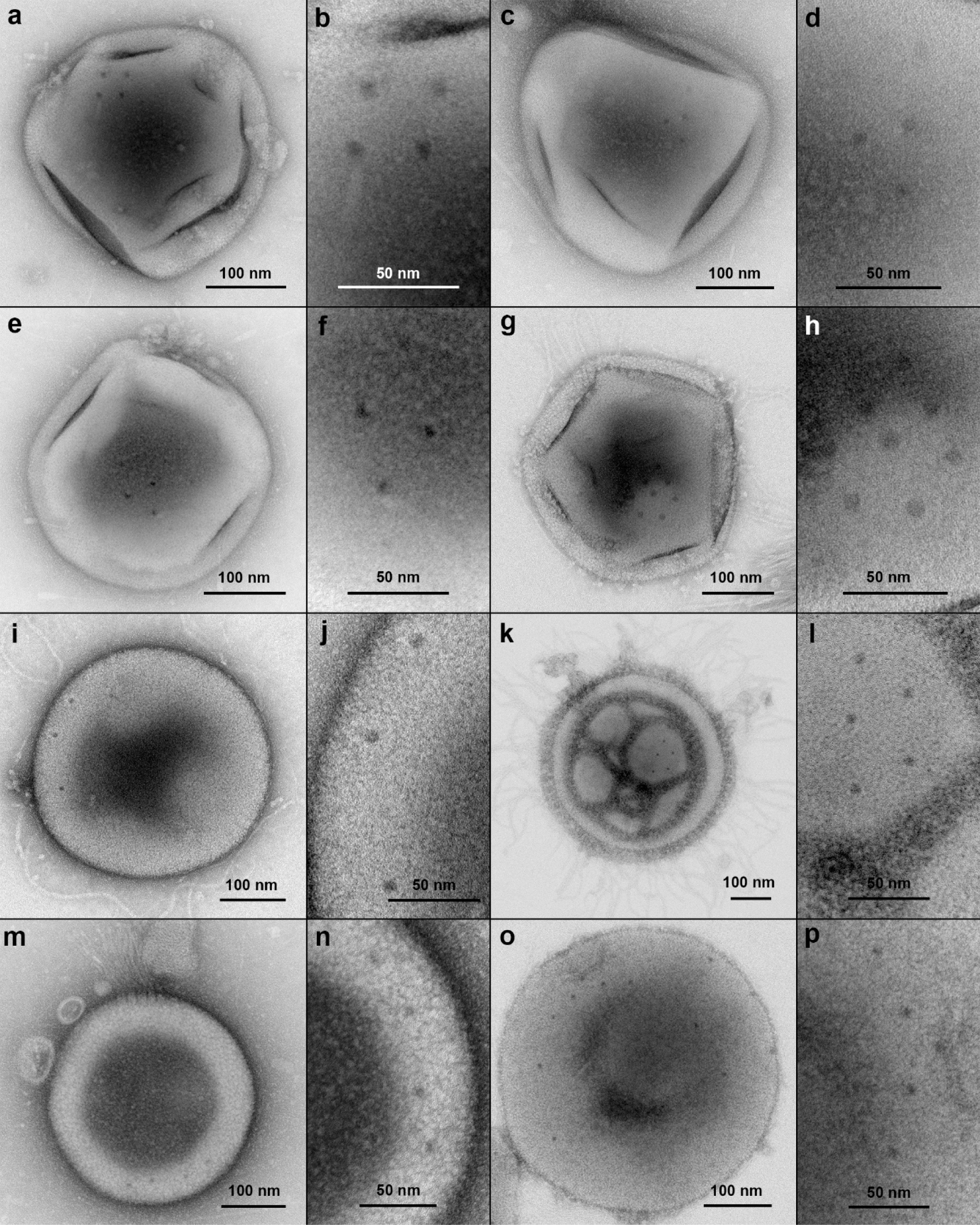
Clustered spots on giant VLPs. Each overview image is accompanied by a high magnification view of the same particle showing arrangements of dark stained pore-like features.

**Supplementary Figure 4:**
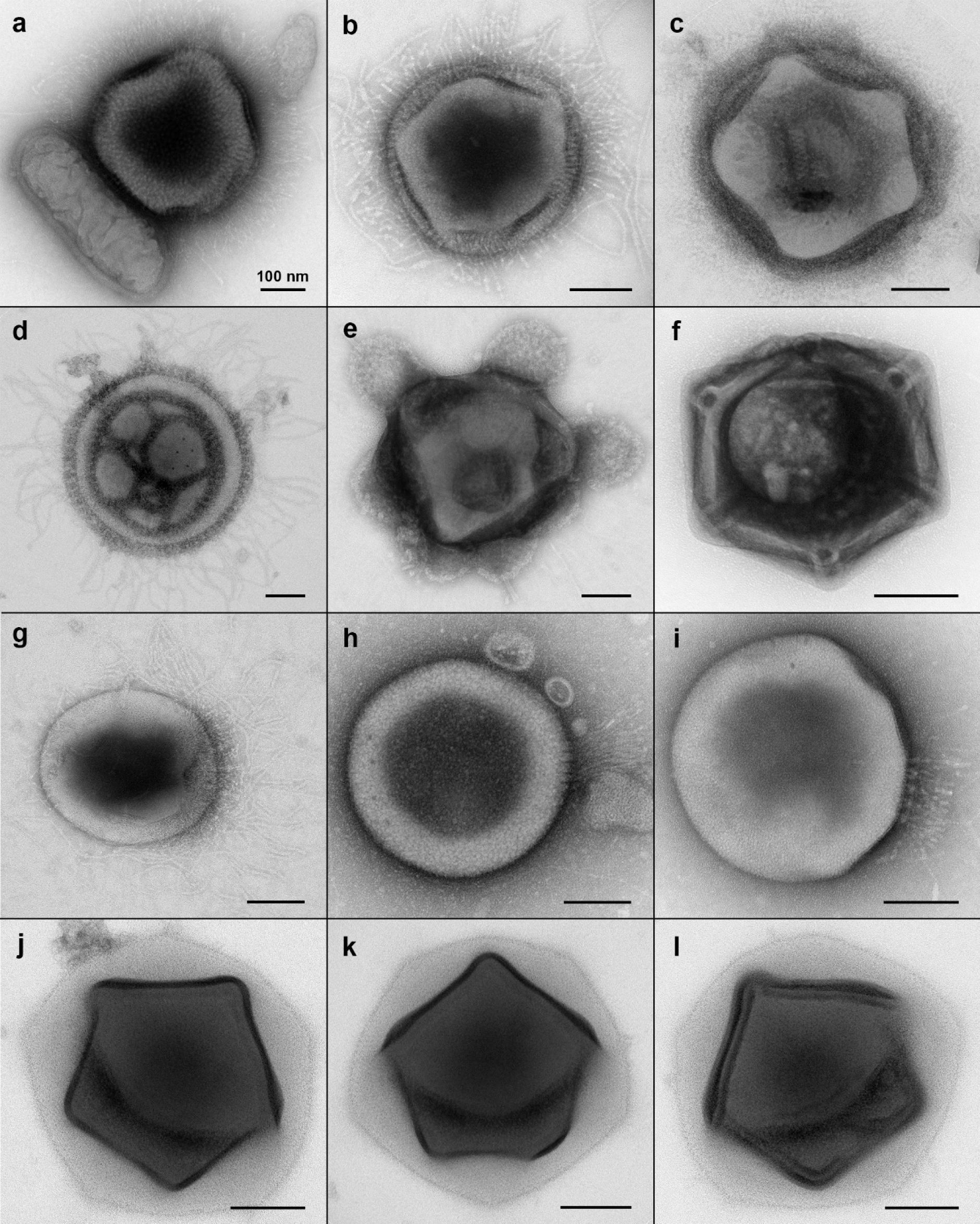
Additional examples of giant VLPs in the categories “mimi-like” (a+b), “supernova”(c+d), “turtle” (e), “plumber” (f), “haircut” (g-i) and “Christmas star” (j-l). All scale bars, 100 nm.

**Supplemental Figure 5:**
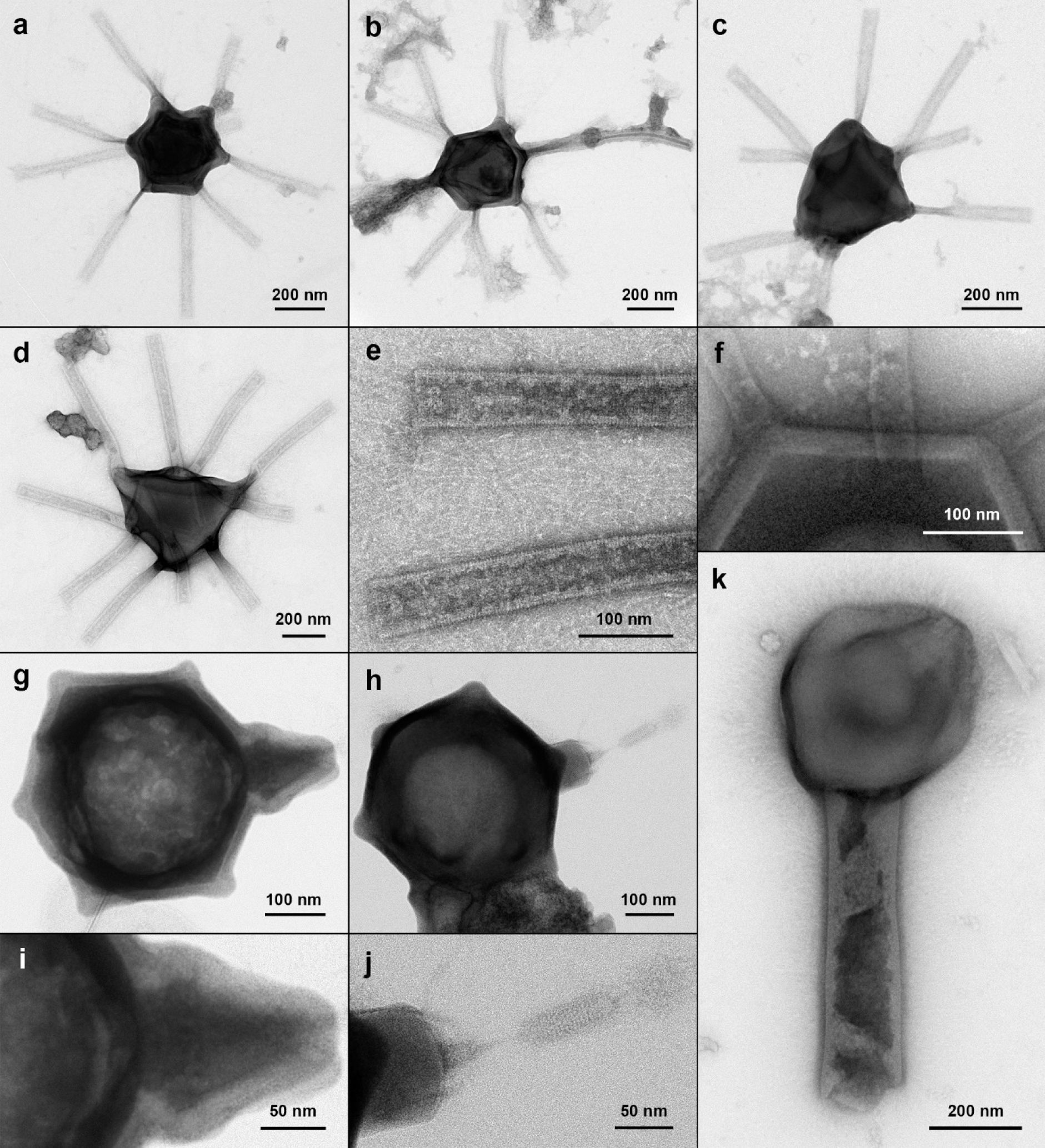
Additional examples of giant VLPs belonging to the morphotypes “Gorgon” (a-d), “flacon” (g+h) and tailed giant VLPs (k). The tubules in e) are a high magnification view of the particle in d); f) is a magnification of Fig. 3h; i) shows a higher magnification of the particle in g); the nozzle structure in j) is magnified from h).

**Supplemental Figure 6:**
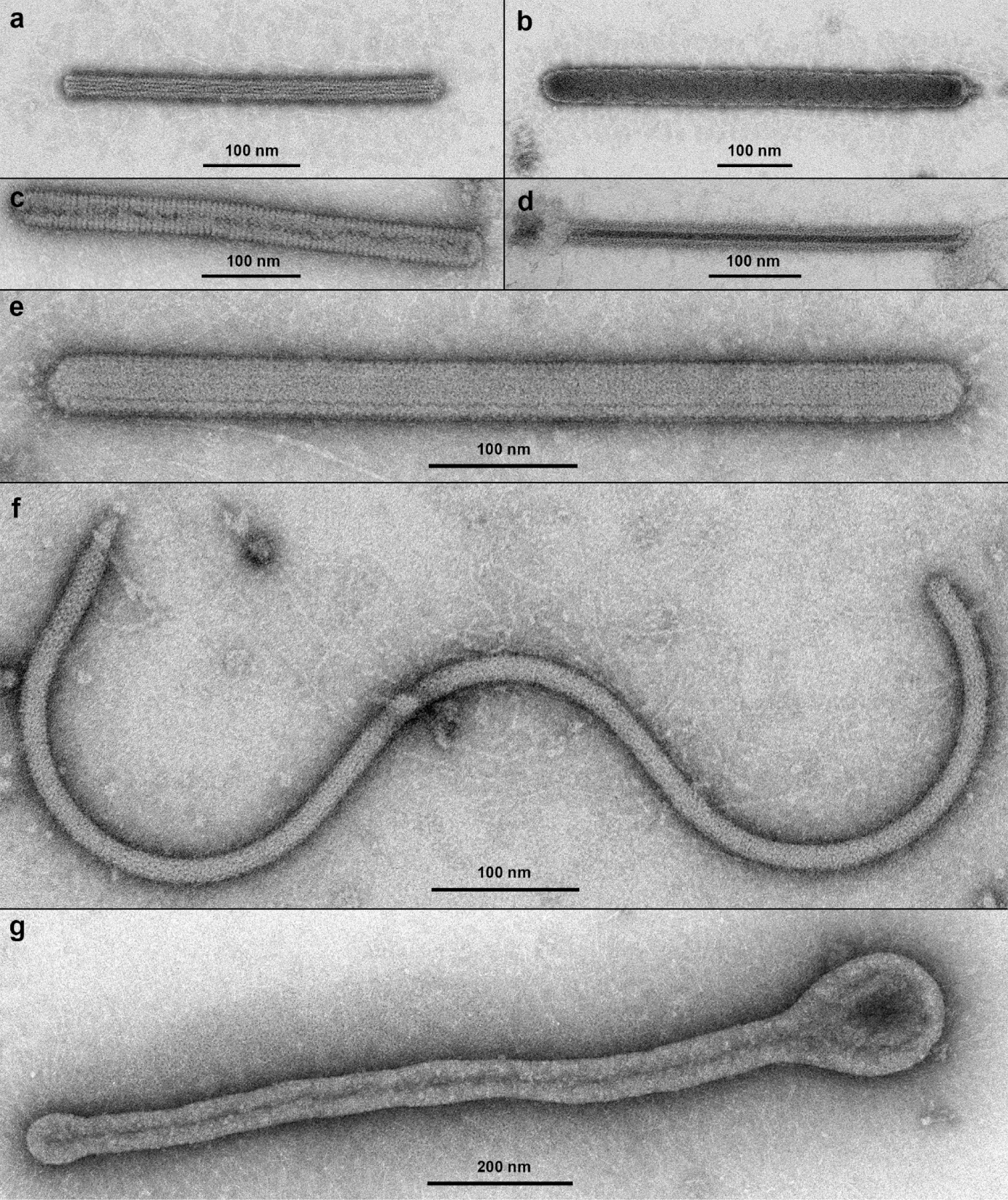
Filamentous virus-like structures. **a)** Rod, 380×20 nm. **b)** Rod, 572×49 nm. This particle could also represent a C3-type podovirus with an extremely elongated head or a B3-type siphovirus with a broken tail. **c)** Rod, 480×36 nm. **d)** Rod, 425×29 nm with an 8 nm-wide electron-dense inner tube. **e)** Rod, 606×38 nm. **f)** VLP or detached cellular structure, 1070 long and 15 nm wide. **g)** Flexible 1270×54 nm VLP with a 165 nm-wide head structure.

**Supplemental Figure 7:**
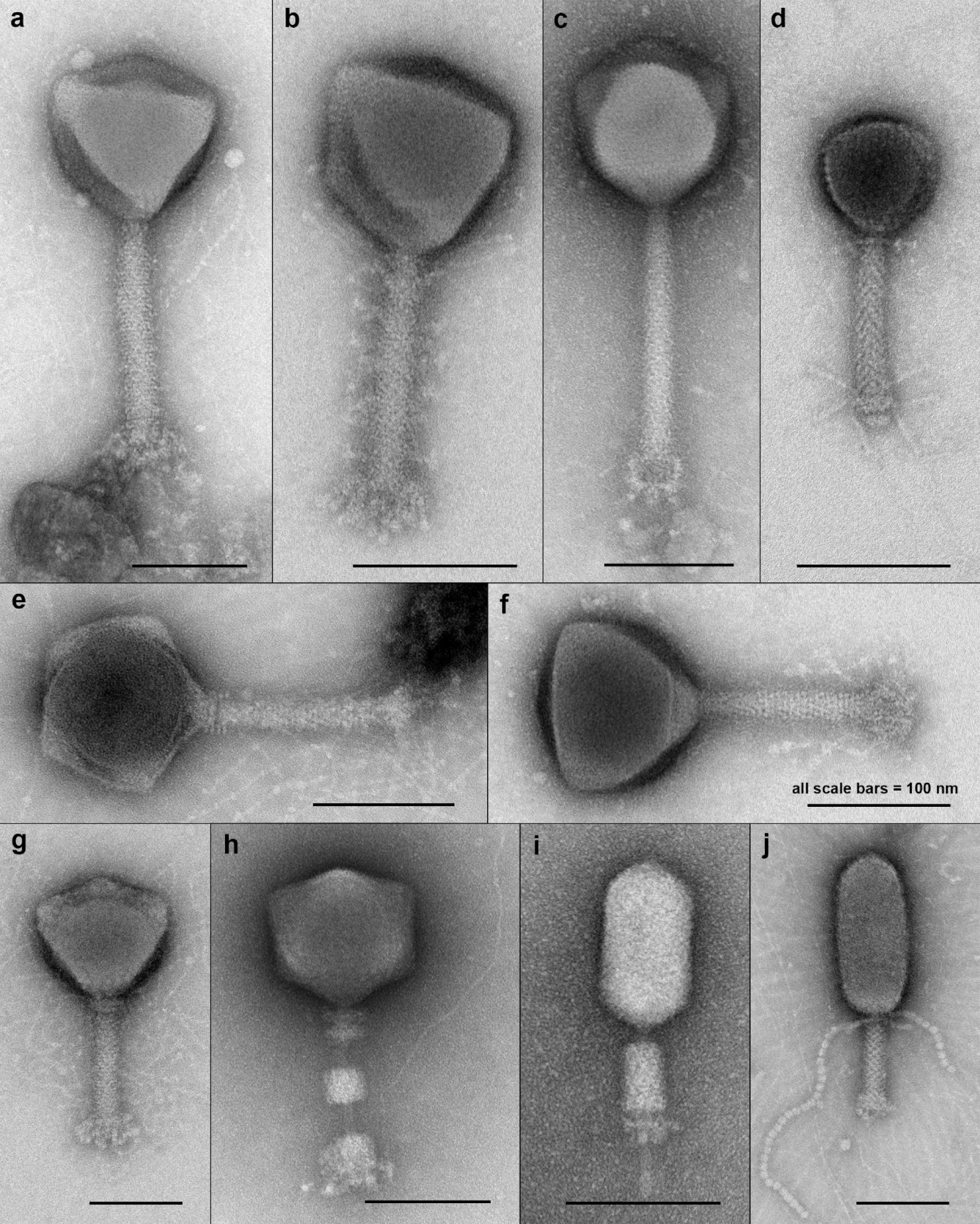
Bacteriophage VLPs with long contractile tails typical of myoviruses. **a)** Head diameter: 150 nm, tail length: 210 nm. **b)** Head diameter: 120 nm, the 168 nm long tail displays associated “whisker-like” fibers with remote resemblance to jumbo coliphage phAPEC6^83^. **c)** Head diameter: 129 nm, tail length: 220 nm. **d)** Head diameter: 75 nm, tail length: 125 nm. **e)** Head diameter: 120 nm, tail length: 145 nm. **f)** Head diameter: 122 nm, tail length: 149 nm. **g)** Head diameter: 135 nm, tail length: 150 nm. **h)** Head diameter: 120 nm, tail length (contracted): 143 nm. **i)** Head dimensions: 105×57 nm, tail length (contracted): 108 nm. **j)** Head dimensions: 170×75 nm, tail length: 100 nm. VLPs in a)-h) correspond to the A1 morphotype, VLPs in i)-j) to the A3 morphotype (according to Ackermann^84^).

**Supplemental Figure 8:**
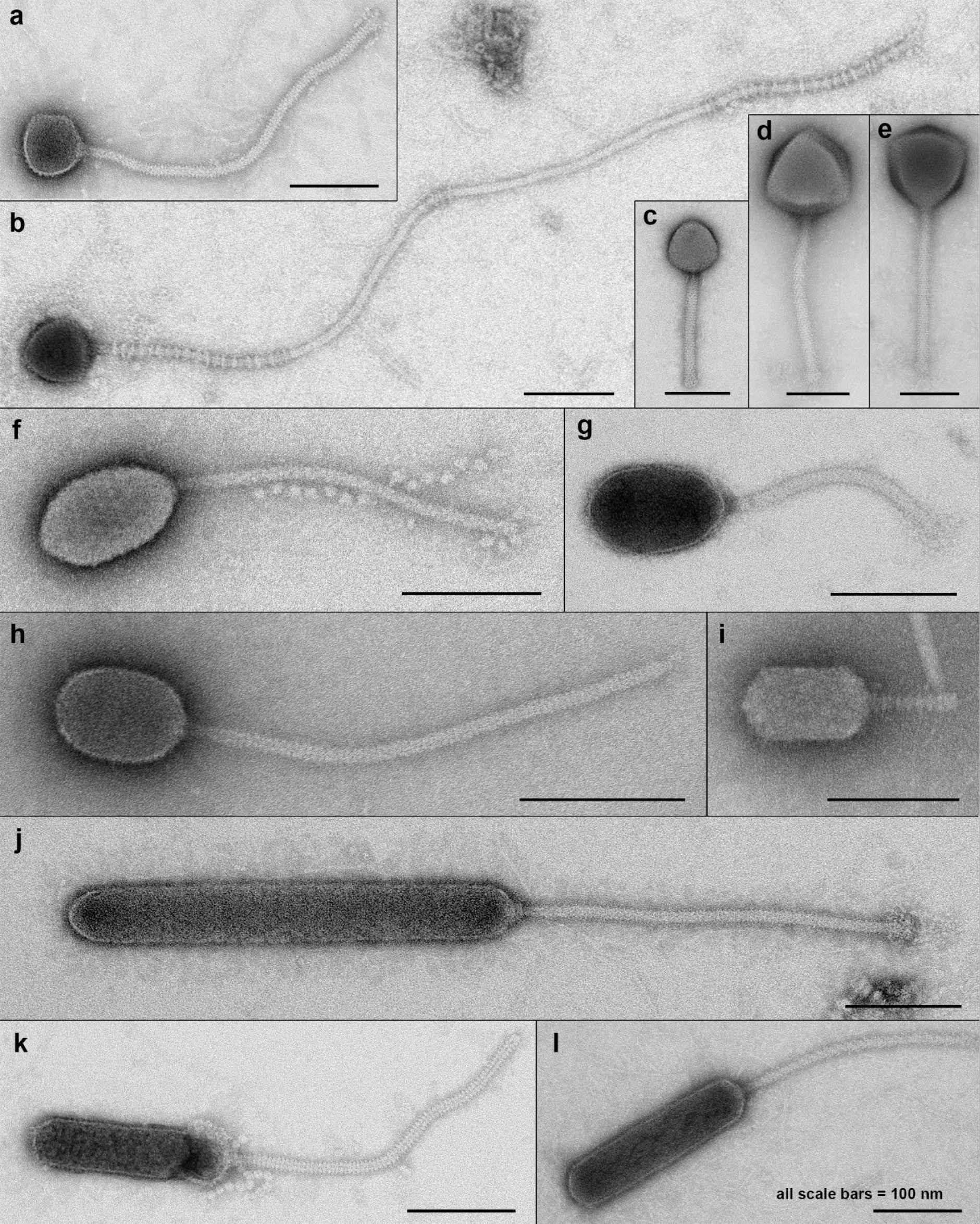
Bacteriophage VLPs with long, non-contractile tails typical of siphoviruses. **a)** Head diameter: 71 nm, tail length: 410 nm. **b)** Head diameter: 71 nm. The tail structure is partially striated and 1.05 µm long. **c)** Head diameter: 77 nm, tail length: 176 nm. **d)** Head diameter: 140 nm, tail length: 268 nm. **e)** Head diameter: 138 nm, tail length: 277 nm. **f)** Head dimensions: 104×62 nm, tail length: 252 nm. **g)** Head dimensions: 111×71 nm, tail length: 190 nm. **h)** Head dimensions: 78×55 nm, tail length: 205 nm. **i)** Head dimensions: 95×54 nm, tail length: 65 nm. **j)** Head dimensions: 387×55 nm, tail length: 348 nm. **k)** Head dimensions: 176×44 nm, tail length: 314 nm. **l)** Head dimensions: 228×56 nm, tail length: 272 nm. VLPs in a)-e) correspond to the B1 morphotype, VLPs in f)-i) to the B2 morphotype, and VLPs in j)-l) to the B3 morphotype (according to Ackermann^84^).

**Supplemental Figure 9:**
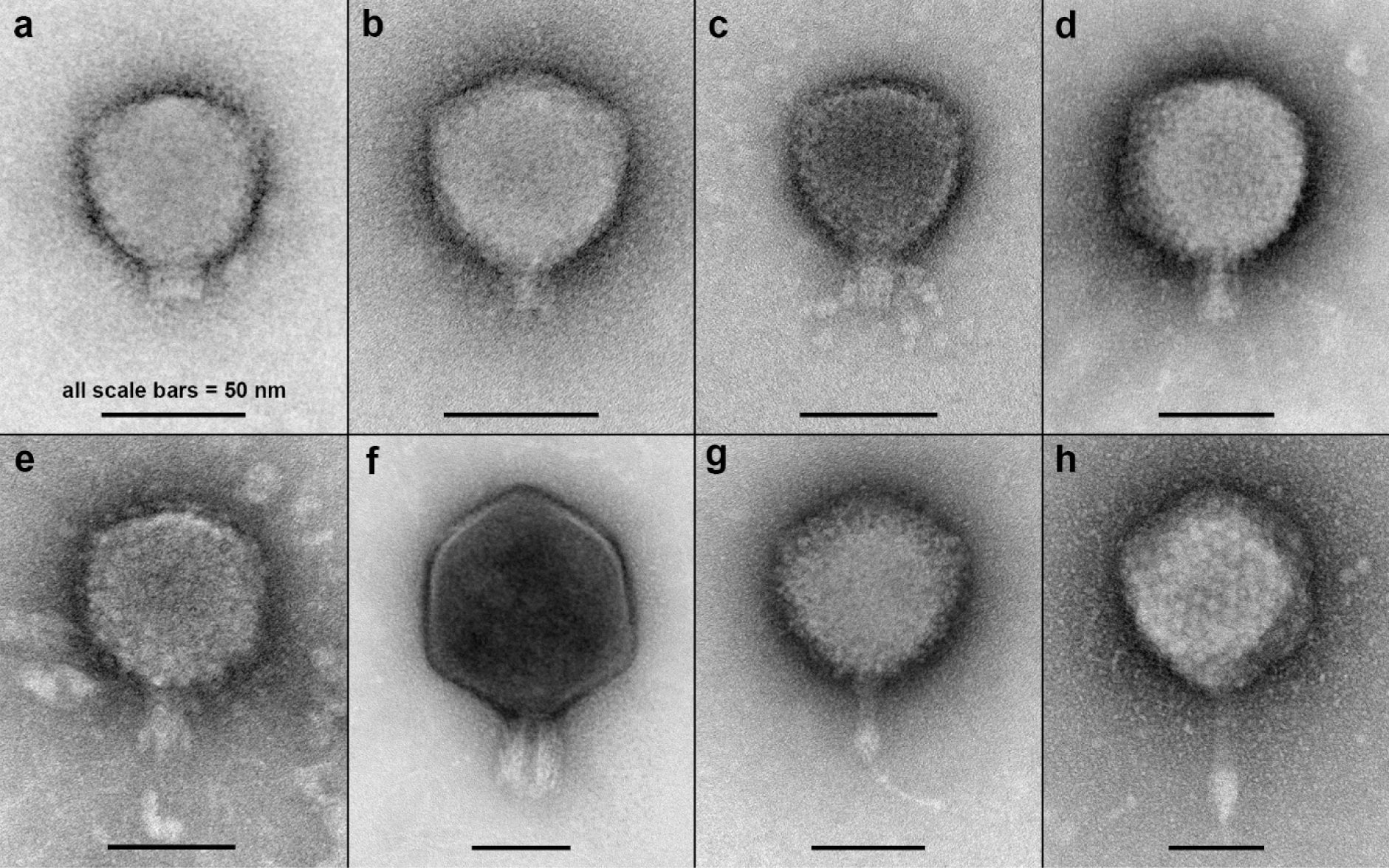
Bacteriophage VLPs with short tails typical of podoviruses. **a)** Head diameter: 62 nm, tail length: 11 nm. **b)** Head diameter: 65 nm, tail length: 13 nm. **c)** Head diameter: 62 nm, tail length: 17 nm. **d)** Head diameter: 85 nm, tail length: 24 nm. **e)** Head diameter: 72 nm, tail length: 22 nm. **f)** Head diameter: 115 nm, tail length: 35 nm. **g)** Head diameter: 83 nm, tail length: 32 nm. A string of material (presumably nucleic acid) is protruding from the tail end. **h)** Head diameter: 107 nm, tail length: 67 nm.

**Supplemental Figure 10:**
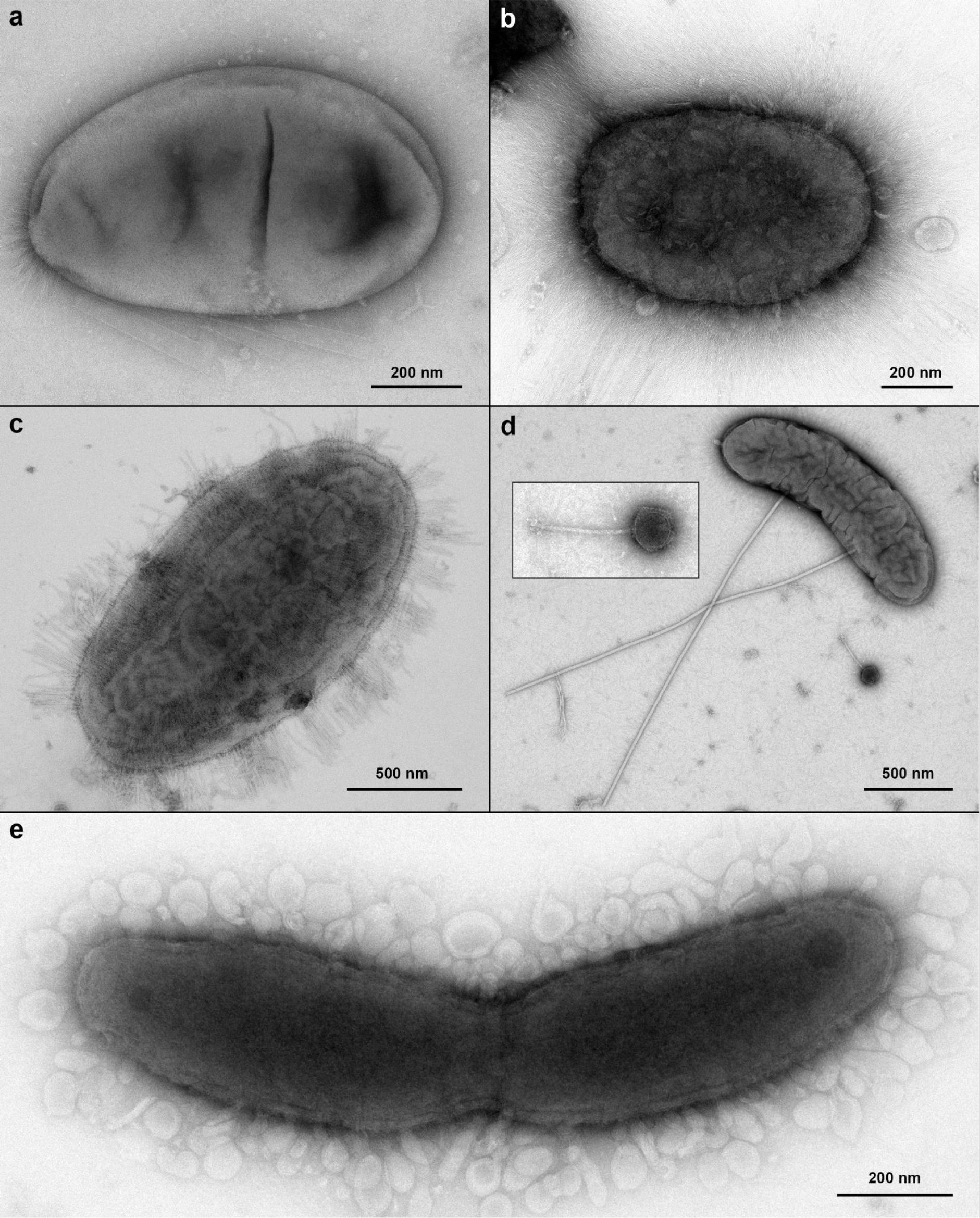
Examples of prokaryote-like particles found in the 0.22 µm - 1.2 µm size fraction of Harvard Forest organic soil. **a)** 940×550 nm b) 830×540 nm with up to 500 nm long hairs. **c)** 1.8×1.0 µm with 200 nm long fimbriae. **d)** 1.5×0.4 µm cell with a pair of 2 µm long, rigid flagellum-like structures. A nearby phage is shown at higher magnification. **e)** 1.45 µm long and 280 nm wide dividing cell with multiple 30-90 nm large vesicle-like structures that interact with the cell surface.

**Supplemental Figure 11:**
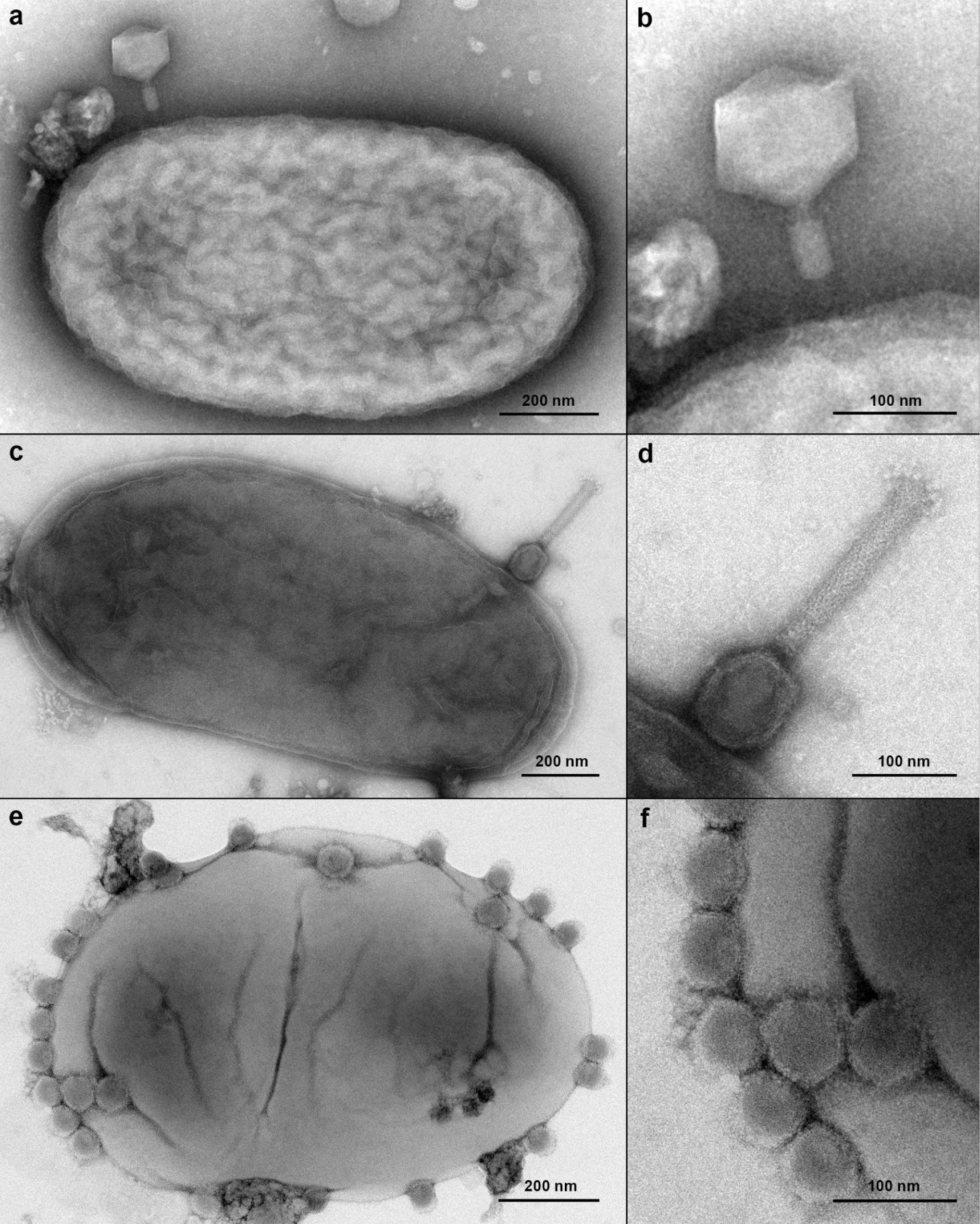
Cells with attached VLPs. **a)**-**b)** Myovirus with contracted tail on bacterial cell surface. **c)**-**d)** Myovirus attached head-first to the cell surface. **e)**-**f)** Multiple VLPs with a diameter of ≈60 nm attached to a cell. Although no tail structures are visible, these VLPs may be podophages.

**Supplemental Figure 12:**
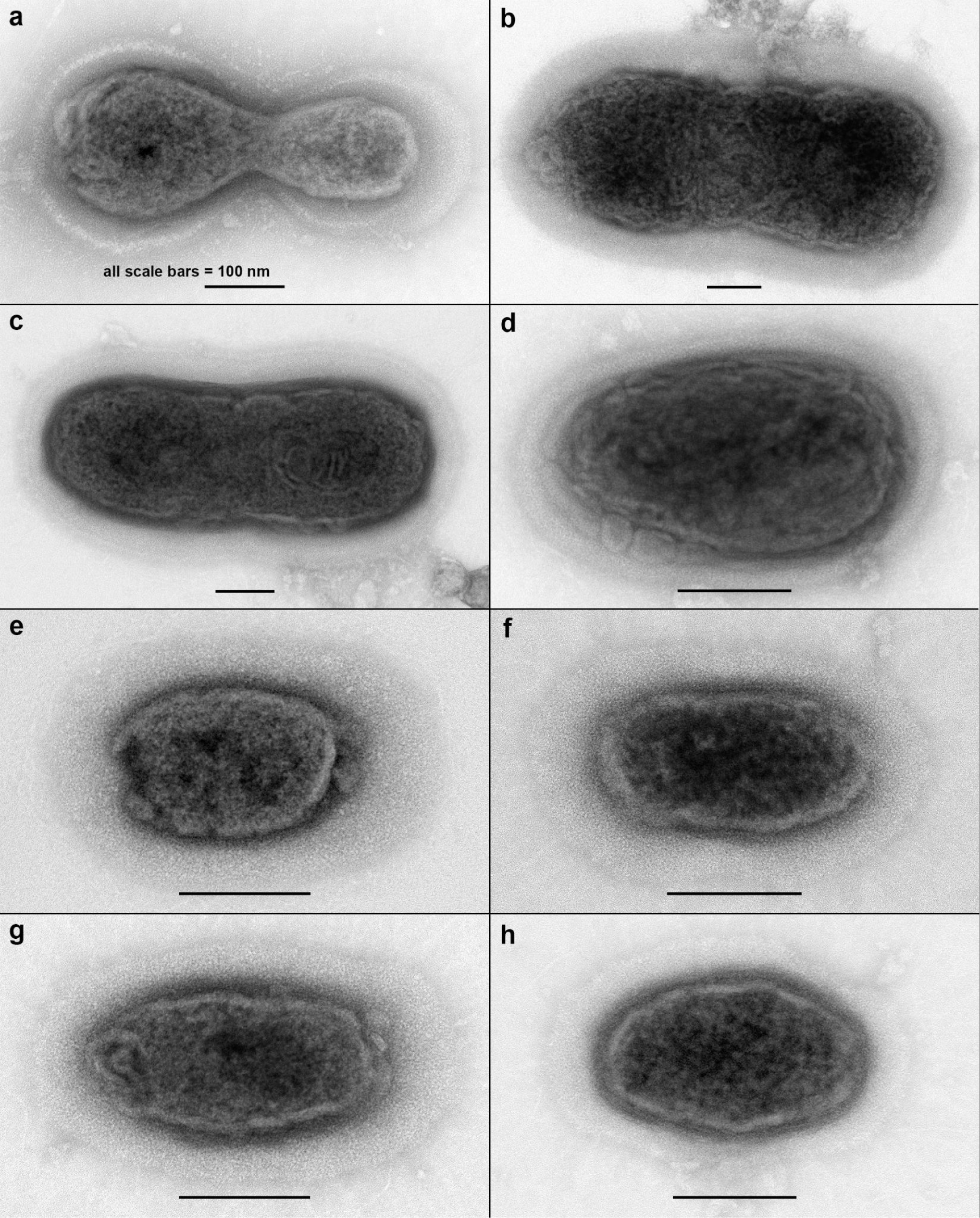
Ultra-small cells with thick surface layers. Dividing cells are shown in a) and b), possibly also in c). The following cell dimensions exclude the outer layer. **a)** 450×180 nm, surface layer: 25-65 nm. **b)** 770×300 nm, surface layer: 47-67 nm. **c)** 675×260 nm, surface layer: 40-55 nm. **d)** 300×180 nm, surface layer: 17-40 nm. **e)** 175×120 nm, surface layer: 30-70 nm. **f)** 200×100 nm, surface layer: 20-45 nm. **g)** 225×100 nm, surface layer: 30-60 nm. **h)** 215×135 nm, surface layer: 20-45 nm. All scale bars, 100 nm.

